# Multiplexed high-content imaging uncovers morphological diversity of lymphocyte activation and dysfunction

**DOI:** 10.64898/2026.02.10.704860

**Authors:** Julie C. Matte, Olivier B. Bakker, Madeline A. Ohl, Francesco Cisternino, Andrea J. Manrique-Rincón, Anke Husmann, Florence Lichou, Tong Li, Kwasi Kwakwa, Anneliese O. Speak, Craig A. Glastonbury, Omer Bayraktar, Carla P. Jones, Melina Claussnitzer, Gosia Trynka

## Abstract

Single-cell transcriptomic and proteomic technologies enable molecular profiling of immune cells at scale but provide limited access to cellular phenotypes shaped by spatial organisation, organelle architecture and cytoskeletal remodelling. Here we present TGlow, a scalable high-content imaging platform optimized for systematic single-cell phenotyping of primary human lymphocytes. TGlow integrates cyclic immunofluorescence, deep z-stack confocal imaging, and open-source data processing pipelines, including both classical and self-supervised vision transformer-based feature extraction, to jointly quantify cellular morphology, organelle organization, and immune activation states. Applied across over 400,000 primary human T cells spanning CD4^+^ activation time courses, drug perturbations, CRISPR knockouts and CD8^+^ T-cell exhaustion, TGlow resolves distinct and reproducible phenotypic states. We uncover dose-dependent and mechanism-specific drug phenotypes, such as defective endoplasmic reticulum polarisation under mycophenolic acid and tofacitinib. We show that mitochondrial clustering reveals activation– and cell-cycle–linked remodelling programs, CRISPR perturbations map gene-specific phenotypes that reposition cells along activation trajectories, and we identify a previously unrecognised collapse of cytoskeletal architecture in exhausted CD8^+^ T cells. TGlow provides a scalable framework for high-dimensional phenotyping of lymphocyte states advancing functional genomics, perturbation screening and population-level immune profiling by resolving the morphological and functional heterogeneity of lymphocytes and enabling systematic linkage of genetic and pharmacological perturbations to cellular function.

## Introduction

Transcriptomic and proteomic profiling have transformed our understanding of cellular function by quantifying RNA and protein abundances at scale and with single cell resolution. Yet, molecular abundance alone does not capture the full spectrum of cellular phenotypes that are shaped by regulatory networks, post-transcriptional and –translational modifications, protein-protein interactions, intracellular trafficking and spatial organisation. Instead, the integrated outcome of these processes often manifests as functional and morphological changes. This is particularly evident in T cell activation, where antigen recognition initiates a cascade of profound transcriptional, metabolic and structural reprogramming that converts previously resting cells into highly motile protein synthesising effector cells. Although omic datasets have characterised the large molecular shifts that occur during these immune processes^1–4^, many represent redundant or indirect effects, obscuring direct mechanistic insights and limiting their interpretability.

Image-based profiling complements omics by capturing morphological and functional phenotypes that are difficult to infer from molecular data alone. High-content imaging (HCI) achieves this through large-scale systematic quantification of thousands of single-cell features captured by fluorescence microscopy^5–7^. Despite its widespread use in drug discovery and functional genomics, HCI and its prevailing assay, Cell Painting, remains underutilized for primary lymphocytes^6–10^. That is because Cell Painting uses a generic, low-cost dye panel that poorly captures immune-specific biology, and the assay was optimized for large, flat adherent cell lines (20–50 µm in xy, ∼5 µm in z), making it technically challenging to apply to small, spherical, non-adherent T cells (5–15 µm in xy and z)^11,12^. While Cell Painting has been adapted for other cell types by substituting dyes with more informative markers, such as a lipid stain in Lipocyte Profiler^6^, comparable adaptations for immune cells have not yet been reported.

There is currently a lack of lymphocyte-oriented HCI assay that jointly multiplexes markers of morphology and immune biological states, capturing features informative of morphology as well as immune activation, proliferation, and metabolic state. To address this gap, we developed TGlow, a scalable image-based profiling assay for phenotypic profiling of lymphocytes at single-cell resolution. TGlow combines cyclic immunofluorescence imaging with classical feature extraction with CellProfiler^13^ or a custom adaptation of the DINO deep learning model^14^ to quantify features of T cell activation (CD25, Ki67), cytoskeleton (actin, tubulin), metabolic indicators (mitochondria, neutral lipids), and other organelles (endoplasmic reticulum (ER), DNA). Core to TGlow is an open-source computational framework, comprising a Nextflow image-processing pipeline (tglow-pipeline; https://github.com/TrynkaLab/tglow-pipeline) and an R analysis toolkit (tglow-r; https://github.com/TrynkaLab/tglow-r), which enables efficient, reproducible single-cell analysis on high-performance computing systems. Here, we apply TGlow to screen CD4^+^ T cell activation timecourses, with drug and CRISPR perturbations, as well as CD8^+^ T cell exhaustion, highlighting its capacity to resolve immune states and uncover biology beyond the reach of standard omics approaches. With its combination of throughput and resolution, TGlow establishes a foundation for unbiased and systematic image-based phenotyping of immune processes, complementing omic approaches to support large-scale projects such as atlasing, perturbation screening or population-level studies to reveal new insights into lymphocyte biology.

## Results

### TGlow: a multiplex high-content imaging ecosystem for profiling T cells

To enable high-throughput and high-content mapping of lymphocyte morphology and activation states, we developed TGlow, a multiplex imaging framework that combines cyclic staining, automated image processing and computational analysis. TGlow performs two sequential imaging cycles to stain eight cellular targets on cells seeded and fixed in 384-well plates (**Fig. 1A, Methods**). Jurkat T cells serve as internal references for inter-plate normalization, mitigating plate-specific variability in signal intensity and enabling objective comparison of features across experimental batches (**Suppl. Fig. 1**). Cycle 1 targets include CD25 (clone M-A251), Ki67 (clone Ki-67), mitochondria (Mitotracker Deep Red), F-actin (actin, Phalloidin) and DNA (Hoechst); and cycle 2, neutral lipids (BODIPY 505/15), ER (ConcanavalinA), tubulin (clone REA1136) and DNA (Hoechst), with parallel brightfield acquisition (**Fig. 1B**). The optical setup (40×, 16 planes with 0.8μm z-spacing; 0.149μm pixel size, 2160×2160 pixels) balances resolution with throughput for small and spherical lymphocytes. Deep sampling in the Z-dimension enables a fair sampling of both resting (∼5μm) and activated T cells (∼12μm), supporting downstream 3D reconstruction and deconvolution, thus enabling an effective resolution improvement of 2.8 times in Z and 1.8 times in X and Y, with a maximal achievable resolution of 0.8μm in Z and 0.24μm in X and Y (**Methods, Suppl. Fig. 2**).

**Figure 1.**
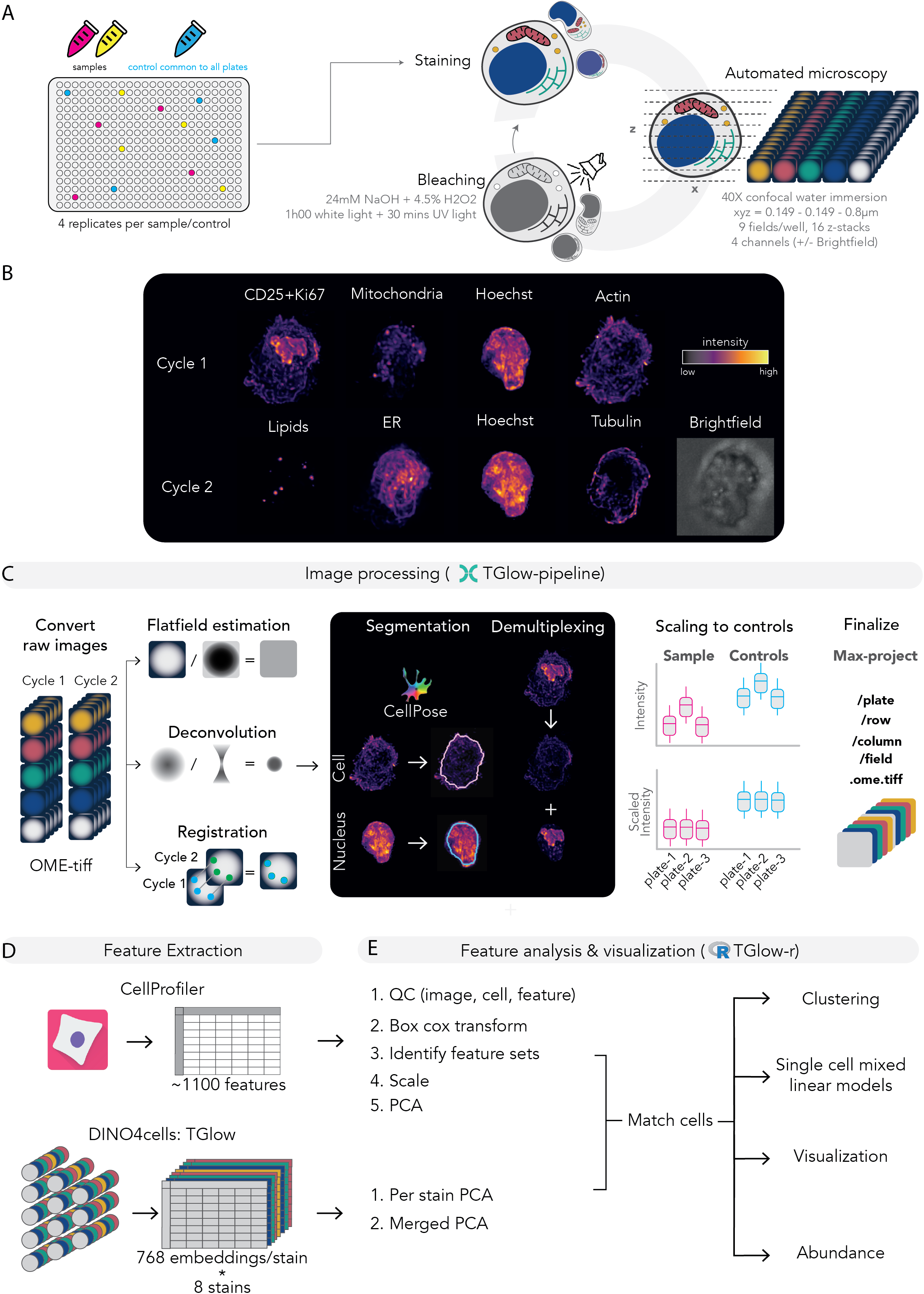
Overview of the TGlow workflow. **A)** Experimental protocol: Samples (yellow and pink tubes) are seeded in quadruplicates onto PLL-coated 384-well imaging plates, excluding the edge wells. Well positions are randomized across the plate and a common control (blue) is included in each plate. Plates undergo two rounds of cyclic imaging in which samples are stained, imaged, chemically (24mM NaOH and 4.5% H_2_0_2_) and photochemically (60 minutes while light and 30 minutes UV light) bleached and restrained for the next cycle. Image acquisition is performed on an Opera Phenix microscope using a 40X water-immersion confocal objective, acquiring 16 z-planes (0.8μm spacing) and nine fields per well. **B)** Representative images (deconvoluted and max projected) of a single cell stained with cycle 1 (CD25+Ki67, mitochondria, Hoechst, actin) and cycle 2 (lipids, ER, Hoechst, tubulin, with brightfield). Stain intensity is displayed from low (black) to high (yellow). **C)** Image Processing through the Nextflow-based TGlow-pipeline. From left to right: microscope exports are converted to OME-TIFF stacks; flatfields, deconvolution and registration are computed in parallel; deconvoluted images are segmented in 3D to delineate the nucleus (Hoechst) and cell boundary (actin); CD25+Ki67 channel is demultiplexed into the nuclear (Ki67) and non-nuclear (CD25) components; plate offsets are calculated based on the common internal control; and finally all of the steps, with the option of max projection, are applied to produce an analysis ready image. **D)** Feature extraction. Morphological descriptors are quantified using CellProfiler (n= 1,098) and/or by generating deep-learning embeddings with a customised DINO4Cells model (768 embeddings for each of the eight channels, 6,144 total). **E)** Feature analysis in R using the *tglow-r* package. CellProfiler features are used for image, cell and feature quality control. CellProfiler features are transformed using BoxCox, optionally scaled and highly correlated features are grouped into ‘feature sets’ based on a feature-feature correlation matrix. For DINO embeddings, the per stain embeddings or an amalgamation of the embeddings of all eight stains (DINO-merged) are subjected to PCA to retain the components that make up 95% of the total variance. Cells of DINO and CellProfiler are mapped to each other by xy position, resolving slight differences in processing. The *tglow-r* package then enables statistical modelling with mixed linear models, clustering with Louvain clustering in PCA space, abundance tests via Milo and visualizations of reductions, features, cluster markers and cells.

Image processing is automated through a customizable Nextflow pipeline designed for scalability on high performance compute clusters (**Fig. 1C**, **Suppl. Fig. 3, Methods**). In short, microscope exports are converted to OME-TIFF stacks and organized in a plate/well/row/field directory structure compatible with filesystems that do not handle large file numbers. The pipeline computes polynomial flatfields, registers imaging cycles using masked cross-correlation, performs Richardson–Lucy deconvolution and segments nuclei and cells in 2D or 3D using CellPose^15^. Deconvoluted and registered images are then flatfield corrected, scaled to internal controls, demultiplexed (e.g., separating nuclear vs non-nuclear markers in the same channel such as CD25 and Ki67) and max projected. Individual cell crops can be exported as HDF5 files compatible with downstream visualization in the *tglow-r* package and training of deep learning models.

TGlow quantifies single cell morphology using CellProfiler and a custom adaptation of DINO4Cells^14^ (**Fig. 1D**). The CellProfiler pipeline extracts 1,098 features, including 1,021 measurements of intensity (e.g., integrated intensity, mean intensity), granularity (i.e., size of globules), morphology (e.g., cell or nucleus size and shape), distribution of intensities (e.g., compactness, radial distribution) and texture measurements (i.e., variances in intensities) for the eight stains across four cellular compartments (nucleus, cytoplasm, membrane, cell) (**Suppl. Fig. 4A, Suppl. Tab. 1)**. The remaining 77 features describe the morphology or network structure of segmented mitochondria, lipids and Ki67. In parallel, DINO4Cells generates 768 dimensional embeddings for each of the eight markers, totalling 6,144 embeddings. To integrate information across all markers, individual DINO embeddings of all eight channels are optionally merged into a single matrix (DINO-merged, **Fig. 1E**).

For statistical analysis, the *tglow-r* package provides a comprehensive R-based environment for single-cell and aggregate-level analyses, including data QC, dimensionality reduction, abundance testing, feature modelling and visualization (**Fig. 1E**). Morphological features are modeled at single-cell resolution using linear mixed-effect models with random effects to account for donor, plate and well variation. Because CellProfiler features often exhibit strong inter-feature correlations, highly correlated features (dissimilarity cut-off of 0.3) are grouped into ‘feature sets’ (**Suppl. Fig. 4B-E, Suppl. Tab. 2, Methods**). Results are summarized at multiple levels: by channel, by channel-category or by feature sets with representative individual features included to illustrate interpretable channel or set-specific properties.

Together, TGlow constitutes an open-source, end-to-end imaging ecosystem that unifies experimental workflows, image processing, and robust statistical analysis to enable scalable, high-content characterization of primary lymphocytes. Although developed for T cells, the framework is broadly adaptable to alternative staining panels and other cell types.

### TGlow captures dose responsive modulation of T cell activation states

To validate TGlow-derived biological inferences, we profiled primary CD4^+^ T cells exposed to drugs targeting cell cycle (palbociclib, hydroxyurea (HU), and mycophenolic acid (MPA)), mitochondrial dynamics (MFI8, mdivi-1, rotenone and disulfiram) and T cell activation (tofacitinib, rotenone). CD4^+^ T cells from three healthy donors were activated for four hours prior to drug exposure, treated with the selected drugs across a range of concentrations and imaged using TGlow at 0h, 24h and 72h post-activation (**Fig. 2A, Methods**). This resulted in 423,010 high-quality single cells across > 350 conditions and 9,748 fields of view underscoring the scalability of TGlow (**Suppl. Fig. 5A)**. The 999 quality-controlled CellProfiler features were grouped into 447 feature sets (**Suppl. Tab. 2**).

**Figure 2.**
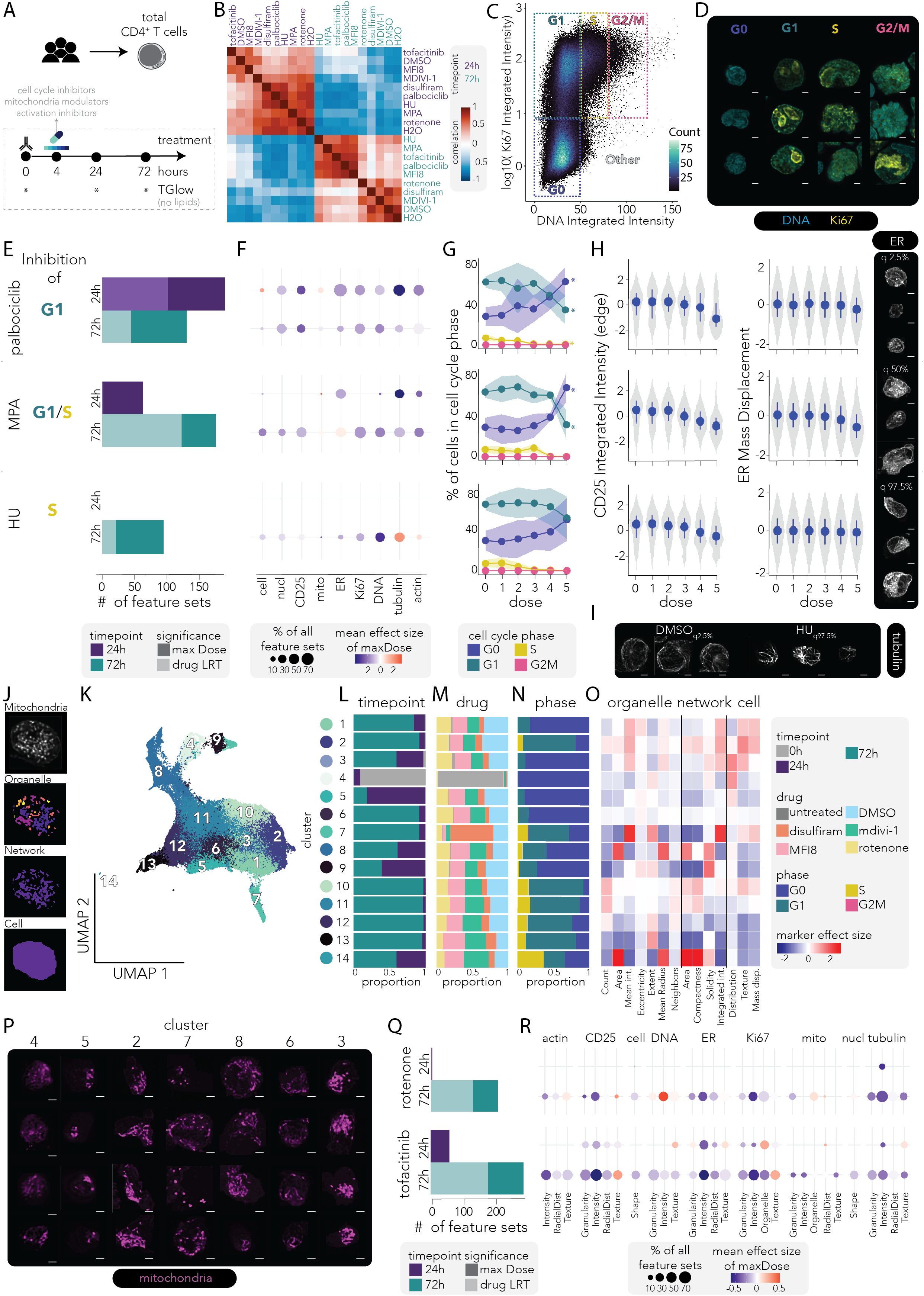
TGlow captures drug-associated dose-dependent morphologies of cell cycle, mitochondrial remodelling and activation. **A)** Experimental design. **B)** Heatmap of pairwise sample profile correlations of the control treatments (DMSO, H2O) and the highest dose of all drugs at 24h and 72h. Features were averaged per sample before correlation. **C)** Cell cycle density dot plot showing log10 (nuclear Ki67 integrated intensity) versus raw nuclear DNA integrated intensity; cell density is coded black to light green (1-100 cells). Dashed outlines indicate manual gates and labels for G0 (blue), G1 (green), S (yellow), G2/M (pink) and Other (grey, label only). **D)** Representative images (deconvoluted and max projected) of each cell cycle phase (randomly selected, n= 3) colored with DNA (blue) and Ki67 (yellow). Scale bar= 2.5μm. **E-H):** Plots relating to the cell cycle drugs: palbociclib (G1 inhibitor, top), MPA (G1/S inhibitor, middle) and HU (S inhibitor, bottom). When applicable, timepoints are shown within a drug: 24h (top) and 72h (bottom). **E)** Barplot depicting the number of differential feature sets per cell cycle drug and timepoint (24h in purple, 72h in blue) modelled as *feature ∼ dose_factor + donor + (1 | plate:well)* and averaged per feature set. Significance either reflects drug term removal (LRT, light shade) or maximum dose (maxDose, dark shade). **F)** Dot plot summarizing significant (maxDose) feature sets by drug/timepoint (rows) and channel (columns, n= 9). Dot size denotes the proportion of significant features in that channel; color encodes mean effect size: –3 (blue) to 0 (white) and +3 (red). **G)** Fraction of cells in each cell cycle phase at 72h (per donor), shown as mean (dot) and SD (ribbon); G0 (blue), G1 (green), S (yellow), G2M (pink). Asterisk (*) indicate significant dose effects (model p≤ 0.05). **H)** Single-cell violin plots of transformed, donor-corrected and scaled values for cell_Intensity_IntegratedIntensityEdge_cd25 (left) and cell_Intensity_MassDisplacement_ER (right). Blue dots and lines represent the median and interquantile range for each dose, respectively. To the right are representative images (deconvoluted and max projected) of the 2.5^th^, 50^th^ and 97.5^th^ quantiles (three each) of cell_Intensity_MassDisplacement_ER, colored by ER (greyscale). Scale bar= 2.5μm. **I)** Representative images (deconvoluted and max projected) of three cells of DMSO treatment (at the 2.5^th^ quantile) and three cells of the HU treatment (at 97.5^th^ quantile) of the feature *cell_Texture_SumAverage_tubulin_12_00_256*, colored by tubulin intensity (greyscale). Scale bar= 2.5μm.**J)**Top to bottom: representative images of the mitochondria stain, organelle segmentation, network segmentation and cell segmentation. **K)** UMAP embedding of 172,898 cells (> 0 mitochondria objects) treated with DMSO, mdivi-1, MFI8, disulfiram or rotenone. Embeddings are derived from 15 PCs calculated from mitochondria features (178 total features: organelle= 38, network= 32 and cell= 108). Cells are colored by cluster annotation (n= 14) and cluster labels are overlaid on the UMAP. **L-O)** Cluster characteristics: **L)** proportion plot of timepoint (0h: grey, 24h: purple, 72h: blue), **M)** proportion plot of drug (untreated: grey, DMSO: blue, disulfiram: orange, mdivi-1: green, MFI8: pink and rotenone: yellow and **N)** proportion plot of cell cycle phase (G0: blue, G1:green, S:yellow, G2M:pink) and **O)** heatmap of cluster marker effect size for 14 representative features spanning organelle (n= 7), network (n= 4) and cell (n= 3). Abbreviations: intensity (int.) and displacement (Disp.) See Suppl. Tab. 4 for full feature names. **P)** Representative images (cell closest to the cluster centroid and its three nearest neighbours, deconvoluted and max-projected) of Clusters 4, 5, 2, 7, 8, 6 and 3 showing mitochondria staining (pink). Scale bar= 2.5μm. **Q)** As described in D), but for rotenone and tofacitinib. **R)** As described in E), but for rotenone and tofacitinib and further separated within channels by category. Mean effect size: –0.5 (blue) to 0 (white) and +0.5 (red).

At the highest doses, sample-sample correlation analyses showed that timepoint was the dominant driver of morphological and functional states (**Fig. 2B**). Increased heterogeneity at 72h indicated that drug-specific phenotypes emerged or intensified over time, with all cell-cycle inhibitors clustering with the activation inhibitor tofacitinib and the mitochondrial inhibitor MFI8, suggesting convergence onto shared downstream phenotypic programs.

Thus, temporal progression primarily shaped CD4^+^ T cell states, with drug perturbations exerting secondary, time-dependent modulatory effects. Overall, drug treatments induced both convergent and divergent feature profiles, with robust and directionally consistent changes across feature sets (**Suppl. Fig 6A-C**). The following sections dissect these responses to validate effects specific to cell cycle alterations, mitochondrial activity and broad T cell activation.

### TGlow captures specific effects of cell cycle inhibitors

Proliferation is a defining effector function of T cell activation and a major source of cellular heterogeneity. As with transcriptomic analyses, cell-cycle effects must be considered in image-based profiling to avoid confounding activation signals while capturing proliferation itself. We therefore tested whether TGlow could detect drug-induced cell-cycle perturbations.

Analogous to flow cytometry cell cycle gating, cells were classified in G0, G1, S, G2M phases based on integrated intensities of DNA and Ki67 (**Fig. 2C**), recapitulating expected phase-specific morphologies (**Fig. 2D**). Without drug treatment, resting T cells were predominantly quiescent (99% G0) and progressed towards more proliferative states following activation: at 24h, 93±1% remained in G0, whereas by 72h, cells shifted to 29 ± 9% G0, 63 ± 6% G1, 7 ± 4% S, and 0 % G2/M (**Suppl. Fig. 5B**).

At 24h, only palbociclib significantly altered cell cycle composition, reducing G1 fractions (β= –1%, p= 0.04), consistent with a ∼30h cell cycle for CD4^+^ T cells (**Suppl. Fig. 5C**). Despite the moderate compositional effects at 24h, the three cell cycle inhibitors induced a number of aberrant morphologies corresponding to their targeted checkpoint (**Fig. 2E**)^16^. At the maximum dose, palbociclib (G1 arrest) induced 190 differential feature sets, MPA (G1/S arrest) 63 and HU (S-phase arrest) 0 (**Suppl. Tab. 3**). When jointly modelling all doses, these numbers decreased to 102, 63 and 0, respectively. Palbociclib effects spanned all channels with the the largest effect sizes observed for tubulin and CD25, whereas MPA particularly perturbed tubulin, actin and the ER: palbociclib impacted 60% of all tubulin feature sets (mean β= –0.4, p_adj-maxDose_≤ 0.04) and 36% of all CD25 feature sets (mean β= –0.3, p_adj-maxDose_≤ 0.05) while MPA impacted 40% of all tubulin feature sets (mean β= –0.4, p_adj-maxDose_≤ 0.05), 20% of actin feature sets (mean β= –0.2, p_adj-maxDose_≤ 0.05) and 53% of ER feature sets (mean β= –0.1, p_adj-maxDose_≤ 0.05, **Fig. 2F, Suppl. Fig. 6B-C**).

At 72h, palbociclib caused a dose-dependent decrease in G1 (β= –14%, p= 6×10^−3^) and S (β= –2%, p= 1×10^−2^) phases, with a parallel increase in G0 (β= 16%, p= 3×10^−3^) (**Fig. 2G, Methods**). MPA induced a dose-responsive reduction in S phase (β= –2.5%, p= 5×10^−3^) and HU caused minor S-phase reductions (β= –0.001%, p= 0.04). Beyond compositional changes, TGlow identified shared and unique sets of morphological and functional signatures for each inhibitor (**Fig. 2H, Suppl. Fig. 6A-C**). While all three impacted T cell activation, indicated by reduced CD25 expression at 72h (R^2^_diff-cell_Intensity_IntegratedIntensity_Edge_= 2%, 4% and 3% with p_adj_= 8×10^−3^, 4×10^−6^ and 1×10^−2^ for palbociclib, MPA and HU, respectively), MPA treatment distinctively perturbed the ER (28% of ER feature sets), with increased intensity but reduced polarization: β_maxDose-cell_Intensity_MeanIntensity_ER_= 1.2 (p_adj_= 1×10^−5^ with R^2^_diff_= 6% and p_adj_= 8×10^−20^) and β_maxDose-cell_Intensity_MassDisplacement_ER_= –0.6 (p_adj_= 2×10^−7^ with R^2^_diff_= 2% and p_adj_= 1×10^−30^, **Fig. 2H**). This implicates ER dysfunction, which is tightly linked to proliferation and cytokine secretion, in MPA’s immunosuppressive mechanism, either as a direct or downstream consequence of its mechanism of action^17–19^.

Furthermore, HU, a ribonucleotide reductase inhibitor that blocks DNA synthesis, induced the characteristic DNA-damage signatures previously observed in T cells^20,21^, but also displayed higher amounts of tubulin with altered textures: HU impacted 46% of DNA feature sets (mean β= –0.3, p_adj-maxDose_≤ 0.03) and 46% of tubulin feature sets (mean β= 0.2, p_adj-maxDose_≤ 0.05). The feature with the largest effect size (β_maxDose-cell_Texture_SumAverage_tubulin_12_00_256_= 0.7, p_adj_= 2×10^−5^) belonged to the feature set *tubulin 18.* Representative images at the extremes of this feature showed a denser tubulin architecture at low values and a more striated architecture at high values (**Fig. 2I**).

HU-induced phenotypes with abundant abnormal elongated tubulin structures have been observed in other cell types and shown to directly cause DNA damage^22^.

Together, these results demonstrate that TGlow sensitively captures dose-dependent effects on both cell cycle composition and morphological or functional states, building deeper mechanistic hypotheses for a drug’s mechanism of action.

### TGlow captures distinct mitochondria morphologies

Mitochondrial remodeling is a hallmark of T cell activation, reflecting shifts in metabolism, signaling, and function^23,24^. Changes in mitochondrial mass, potential and network structure define activation states, but are challenging to measure at scale because mitochondria are small and lymphocytes compact, requiring high-resolution imaging^25^. We therefore benchmarked TGlow-derived mitochondrial phenotypes against known biology and drug responses.

To map mitochondria remodelling during T cell activation, we treated cells with disulfiram (acetaldehyde dehydrogenases 2 inhibitor), mdivi-1 (fission inhibitor), MFI8 (fusion inhibitor) and rotenone (mitochondrial complex 1 inhibitor). High doses of mdivi-1 (50**μM** – 100**μM**), MFI8 (100**μM**) and disulfiram (10**μM**) increased G0 proportions by 72h, indicating potential mitochondrial toxicity (**Suppl. Fig. 5C**). To characterize the morphological diversity of mitochondria, we applied unsupervised clustering to 178 CellProfiler-derived mitochondria features describing organelle (n= 38), network (n= 32) or cell-level (n= 108) morphologies (**Fig. 2J**), resolving 14 distinct clusters (**Fig. 2K**). These mitochondrial clusters reflected time-, drug– and cell cycle-dependent events (**Fig. 2L-N**, respectively); and were defined by representative features such as mitochondrial count, size, elongation and activity or network size, fullness, cellular distribution and polarization (**Fig. 2O-P, Suppl. Tab. 4**).

Resting T cells predominantly occupied Cluster 4 (91% of Cluster 4 are resting; 80% of resting cells are in Cluster 4), characterized by low-intensity, centrally distributed mitochondria in a full network, consistent with the reduced oxidative phosphorylation state of resting T cells^26–30^: β_mito_Intensity_MeanIntensity_mito_= –0.4 (p_adj_= 8×10^−243^), β_cell_RadialDistribution_FractAtD_mito_1of6_= 0.9 (p_adj_< 3×10^−323^), β_mitoNetwork_AreaShape_Solidity_= 0.1 (p_adj_= 10×10^−5^). Upon activation, mitochondrial morphology diversified, giving rise to clusters associated with timepoint and cell cycle phase. For instance, Cluster 5 displayed a small network of slightly elongated mitochondria and was enriched for 24h-G0 cells (81%): β_mitoNetwork_AreaShape_EquivalentDiameter_= –0.9 (p_adj_< 3×10^−323^), β_mito_AreaShape_Eccentricity_= 0.2 (p_adj_= 9×10^−25^). This may reflect relocalization of the mitochondria to the synapse before entry into the cell cycle^31^. Conversely, Cluster 2 comprised 72h-G1 cells (91%) featuring numerous large and highly active mitochondria in an expanded network, consistent with elevated energetic demands during proliferation^23^: β_cell_Children_mito_count_= 0.7, β_mito_AreaShapeArea_= 0.3, β_mito_Intensity_MeanIntensity_mito_= 0.8 and β_mitoNetwork_AreaShape_EquivalentDiameter_= 0.5 (all with p_adj_< 3×10^−323^).

Drug treatments produced distinct remodeling signatures. Disulfiram enriched Cluster 7 (60% of Cluster 7 are 72h-disulfiram cells; 30±15% of 72h-disulfiram cells are in Cluster 7), characterized by an intense mitochondria signal (β_mito_Intensity_MeanIntensity_mito_= 2.0 and p_adj_< 3×10^−323^), consistent with disulfiram-induced protein instability leading to an accumulation of protein aggregates in the mitochondrial matrix^32^. Low doses of mdivi-1, a fission inhibitor, increased the proportion of Cluster 8 from 3±2% to 20±8% (R^2^_diff_= 68%, p= 0.02) which featured the largest mitochondria in the dataset (β_mito_AreaShapeArea_= 1.8, p_adj_< 3×10^−323^), and Cluster 6 from 7±1 to 11±2% (R^2^_diff_= 60%, p= 0.05) which displayed elongated mitochondria (β_mito_AreaShape_Eccentricity_= 0.2, p_adj_= 3×10^−60^). Mdivi-1 treatment also led to an overall increase in the size of the mitochondria network, albeit with reduced intensity, consistent with its role as a fission inhibitor: R_2diff-mitoNetwork_AreaShape_MajorAxisLength_= 1% (p_adj_= 5×10^−4^) and R^2^_diff-mito_Intensity_MedianIntensity_mito_= 9% (p_adj_= 1×10^−2^; **Suppl. Tab. 3**). Conversely, inhibition of fusion with MFI8 at the highest dose increased the proportions of cells in Clusters 1, 4 and 5: R^2^_diff_= 77%, 91% and 66% (p≤ 0.03), respectively. These clusters were mostly composed of G0 cells (83%, 99%, 90%, respectively), possibly indicating a toxic effect pushing the cells into quiescence. Finally, rotenone treatment had modest effects on mitochondria morphology with an overall decrease in mitochondria granularity: R^2^ 1% (p_adj_= 1×10^−5^, **Suppl. Tab. 3**).

These results highlight the tight coupling of mitochondria architecture and function and demonstrate that TGlow robustly maps mitochondria morphology, capturing heterogeneity driven by activation, cell cycle and drug perturbation.

### TGlow maps mechanism-specific drug responses of T cell activation

A central motivation of high-content profiling is to capture how mechanistically distinct perturbations reshape cell morphology and function to infer mechanisms of action. To test whether TGlow can differentiate such pathways during activation, we examined inhibitors with contrasting mechanisms: rotenone, which disrupts mitochondrial complex I and cellular energetics essential for activation, cytokine production and proliferation^33^, and tofacitinib, which inhibits JAK-STAT signaling, thereby dampening cytokine-driven activation cues such as IL-2 and IFNγ that are essential for Th1 priming, T-bet induction and early effector differentiation ^34–36^.

Rotenone showed a delayed impact evident only at 72h whereas tofacitinib induced early morphological and metabolic changes, consistent with their respective roles as a cumulative metabolic inhibitor and an activation signalling inhibitor (**Fig. 2Q-R**): at 24h, tofacitinib altered 12% of feature sets compared with 1% for rotenone (0.5% and 1% dose-dependent, respectively) and by 72h, these rose to 57% and 41% (35% and 25% dose-dependent). The early tofacitinib response at 24h involved canonical markers of activation, including a reduction in CD25 expression, a reduction in proliferation and an increase in DNA texture, suggesting chromatin condensation following JAK/STAT blockade and thus reduced transcriptional activity^37–40^: impacted were 44% of all CD25_intensity (mean β= –0.2, p_adj-maxDose_≤ 0.04), 12% of Ki67_intensity (mean β= –0.3, p_adj-maxDose_≤ 0.04) and 29% of DNA_texture (β= 0.2, p_adj-maxDose_≤ 0.04) feature sets. The latter is exemplified by *nucl_Texture_Correlation_dna_12_00_256* (β= 0.1, p_adj_= 0.04, *DNA_texture 3*) and *nucl_Texture_InfoMeas2_dna_3_00_256* (β= 0.2, p_adj_= 0.02, *DNA_texture 4*).

Additional dysregulated phenotypes included altered distribution of the mitochondria, reduced tubulin abundance, and broad suppression of ER-associated features, with the largest effects on ER abundance: impacted were 4% of all mito_RadialDistribution feature sets (mean β= 0.2, p_adj-maxDose_≤ 0.04), 17% of all tubulin_Intensity feature sets (mean β= –0.4, p_adj-maxDose_≤ 0.05) and 62% of all ER_Intensity feature sets (mean β= –0.4, p_adj-maxDose_≤ 0.05). Tofacitinib-induced dysregulation of the ER compartment persisted through 72h, characterized by both a marked reduction in ER abundance and altered intra-cellular organisation: impacted were 75% of all ER_Intensity feature sets (mean β= –0.6, p_adj-maxDose_≤ 0.01) and 55% of all ER_RadialDistribution feature sets (mean β= –0.2, p_adj-maxDose_≤ 6×10^−3^). A representative feature of the feature set ER_Intensity 5 is *cell_Intensity_MassDisplacement_ER* (β= –0.4, p_adj_= 3×10^−15^). Given that tofacitinib decreases the frequency of IFN-producing cells^36,41^, this data suggest that modulation of ER content and organization that is so crucial for cytokine secretion may contribute to this phenotype.

In contrast, the feature sets impacted by rotenone in early activation were confined to tubulin intensity and cell shape: 25% of all tubulin_Intensity feature sets (mean β= –0.5, p_adj-maxDose_≤ 0.04) and 2% of all cell_Shape feature sets (mean β= –0.2, p_adj-maxDose_≤ 6×10^−3^). By 72h, rotenone exposure generated a distinct profile of T cell suppression with the affected feature sets distributed across all compartments, including the expected mitochondrial remodelling^33^. Many of these phenotypes recapitulated cellular processes observed in other cell types but not previously described in T cells, including tubulin dysregulation, actin instability and ER stress response^42–46^: impacted were 92% of tubulin_intensity (mean R^2^ = 7%, p_adj_≤ 1×10^−4^), 57% of tubulin_RadialDistribution (mean R^2^ = 4%, p_adj_≤ 6×10^−5^), 25% of actin_intensity (mean R^2^ = 4%, p_adj_≤ 1×10^−4^), 33% of actin_texture (mean R^2^ = 2%, p_adj_≤ 8×10^−5^), 62% of ER_texture (mean R^2^ = 3%, p_adj_≤ 2×10^−5^) and 75% of ER_intensity (mean R^2^ = 3%, p_adj_≤ 1×10^−5^) feature sets. Together, these data demonstrate that TGlow sensitively resolves the temporally and mechanistically distinct morphological responses induced by metabolic and signalling inhibitors, generating profiles of T cell suppression that recapitulate known biology.

### TGlow uncovers diverse activation states across CD4^+^ T cell populations

Having comprehensively benchmarked TGlow, we applied it to explore a fundamental immunological question: *how do T cell morphological and functional states evolve during activation*? To do so, we profiled naive and memory CD4^+^ T cells from four donors in their resting state (0h) and at four timepoints after anti-CD3/CD28/CD2 stimulation (12h, 2 days, 5 days and 7 days, **Fig. 3A; Methods**). After QC, we recovered 11,749 cells described by 6,144 DINO features and 465 CellProfiler feature sets (**Suppl. Fig. 7A, Suppl. Tab. 2**). We selected these two populations of CD4^+^ T cells as they are very similar in morphology. However, in comparison to memory cells, naive cells have previously not undergone activation, therefore, the response to activation between the two cell subsets is different. We examined whether TGlow could resolve shared and population-specific morphological and functional phenotypes of activation.

**Figure 3.**
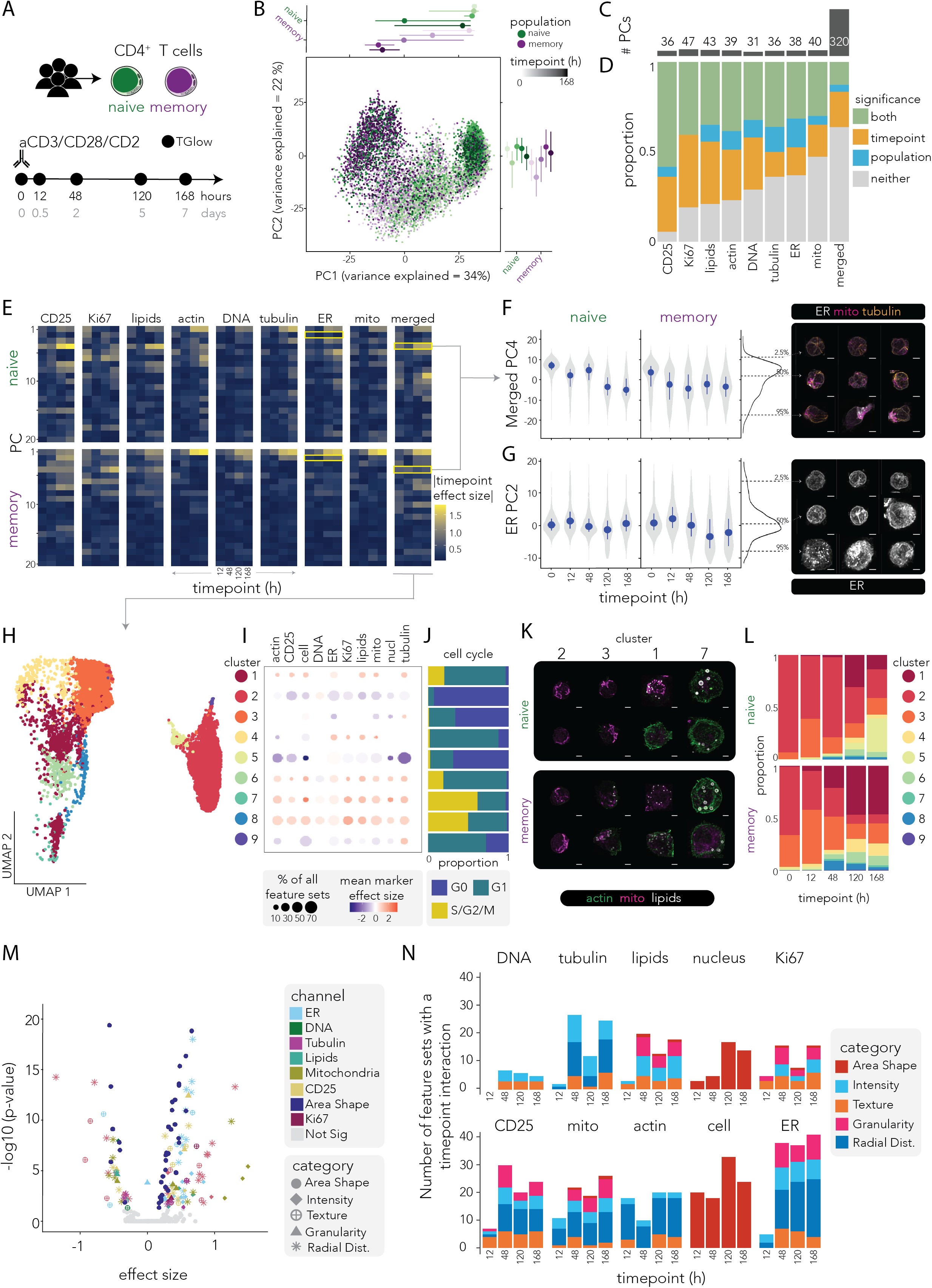
TGlow, combined with deep-learning features, captures morphologies associated with T cell activation. **A)** Experimental design. **B)** PCA calculated from the merged DINO embeddings (n= 6,144). Center: Dotplot of PC1 (variance explained= 34%) and PC2 (variance explained= 22%) with single cells colored by population (naive: green, memory: purple) and shaded by timepoint (0h to 168h in increasing darkness). Below (PC1) and to the left (PC2) are line plots summarising the median and interquantile ranges of each timepoint:population combination for the respective PCs, with the same color and shading code. **C)** Barplot showing the number of DINO-PCs that explain 95% of the total variance in individual channel (CD25, Ki67, lipids, actin, DNA, tubulin, ER, mito) or in the combination of all DINO-embeddings (merged). **D)** Proportion plots depicting the fraction of the selected DINO-PCS in C) are significantly associated with timepoint (orange), population (blue), both (green) or neither (grey) after LRT and multiple testing correction. Full model: *feature ∼ timepoint*population + donor + (1|plate:well)*, Reduced models for timepoint: *feature ∼ population + donor + (1|plate:well)* and population: *feature ∼ timepoint + donor + (1|plate:well).* **E)** Heatmap facetted by channel (columns, same order as D) and by population (row, naive: top, memory: bottom) showing the first 20 PCs (y-axis) by timepoint (12-168h, x-axis); and filled by absolute effect size of timepoint relative to 0h (blue to yellow, 0 to 2). Yellow boxes highlight the PCs shown in F) and G). **F-G)** Violin plots of the scaled merged-PC4 (**panel F**) and ER-PC2 (panel G) over timepoint and separated by column in naive and memory populations. To the right is a histogram of the respective PCs with dotted lines representing the 2.5%, 50% and 97.5% quantiles, followed by representative images (n= 3 per quantile, deconvoluted and max projected). Image colors: ER (white), mito (pink, only in panel F), tubulin (orange, only in panel G) Scale bar= 2.5μm. **H)** UMAP embedding of the merged assay where each cell is colored by a cluster annotation (n= 9, resolution= 0.4, nPCs= 30). **I)** Marker plot where the size of the dot represents the percentage of significant feature sets per cluster (row) in each channel (column) and the color depicts the mean marker effect size from –3 (blue) to 0 (white) and +3 (red). **J)** Proportion per cluster (row) filled by cell cycle phase: G0 (blue), G1 (green), S/G2/M (yellow). **K)** Representative images (n= 2 per population per cluster, cell closest to the centroid and its neighbour, deconvoluted and max projected) of Clusters 2, 3, 1 and 7 with mitochondria (pink), actin (green) and lipids (white). Scale bar= 2.5μm. **L)** Proportion plot per timepoint (x-axis) and population (naive: top row, memory: bottom row) filled by cluster annotation (n= 9, same coloring as H-I)). **M)** Volcano plot displaying the mean main effect size of the feature sets (x-axis) versus the –log10(p-adjusted) from modelling *feature ∼ timepoint*population + donor + (1|plate:well)* and averaged per feature set (n= 430). Naive cells are set as a reference population and timepoints are set as factors. The main effect size can be interpreted as the difference in naive and memory CD4^+^ T cells at resting. Each dot represents the mean of a feature set, with grey circles indicating no statistical dysregulation and otherwise, colors indicating channels (ER: light blue, DNA: green, tubulin: purple, lipids: teal, mitochondria: green, CD25: yellow, AreaShape: dark blue, Ki67: burgundy) and shapes indicating categories (AreaShape: large circle, Intensity: diamond, Texture: circle and cross, Granularity: triangle, Radial Distribution (Radial Dist.): star). **N)** Barplot of the number of feature sets with a significant interaction terms (p_adj_ < 0.05) per timepoint, faceted by channel and colored by category (AreaShape: red, Intensity: light blue, Texture: orange, Granularity:pink, RadialDistribution: dark blue).

### Deep-learning embeddings reveal activation and population-specific morphodynamics

To assess whether DINO embeddings captured patterns linked to T cell activation and population, we applied variance decomposition to the DINO-merged assay (combination of all channel embeddings). After accounting for all covariates, most variation was explained by cell cycle (R^2^_median_= 15%, R^2^_max_= 64%) and timepoint (R^2^_median_= 6%, R^2^_max_= 35%, **Suppl. Fig. 7C-D**), confirming that morphological diversity is largely driven by proliferation and temporal dynamics. Population, its interaction with timepoint, and donor identity accounted for smaller, but detectable fractions of variance: R^2^_median-population_= 1%, R^2^_max-population_= 9%, R^2^_median-population:timepoint_= 2%, R^2^_max-population:timepoint_ = 18%, R^2^_median-donor_ = 0.1%, R^2^_max-donor_ = 1%.

PCA of the DINO-merged assay showed separation of naive and memory cells across timepoints along PC1 (34% of the total variance explained) and PC2 (22%, **Fig. 3B**). To isolate channel-specific information and reduce dimensionality for downstream analyses, we identified the number of PCs required to explain 95% of the variance in the DINO-merged and the individual channel embeddings. The merged assay required 320 PCs while the single-channel embeddings required 31 to 47 PCs (DNA= 31, CD25= 36, tubulin= 36, ER= 38, actin= 39, mitochondria= 40 lipids= 43, Ki67= 47, **Fig. 3C**). These differences indicate that channels differ in the complexity of morphological variation they encode, whilst the similar overall dimensionality of the merged assay and the sum of the individual channels (n= 310) implies non-redundant information across channels.

To quantitatively assess differences associated with activation and population, we modeled DINO PCs of the individual channels and the merged channels using a linear regression framework that included an interaction between population (naive or memory) and timepoint (treated as a factor) (**Fig. 3D-E, Suppl. Tab. 5**). A quarter of PCs showed effects for both timepoint and population, capturing morphological and functional programs that differed between naive and memory cells while evolving with activation. Timepoint alone explained 23% of PCs, capturing conserved activation-driven changes, while population alone accounted for 6%, reflecting differences between naive and memory cells that are stable across activation. Among channels, CD25 and Ki67 showed the highest proportion of timepoint-associated PCs (94% and 81%, respectively), consistent with their roles as canonical markers of activation and proliferation. Whereas Ki67-PCs showed no population-specific effects, which is expected given proliferation is a cell type agnostic process, 5% (n= 2) of the CD25-PCs showed population-specific effects, reflecting the higher baseline expression in memory cells^47^. The ER (16%, n= 6), tubulin (14%, n= 5), lipids (11%, n= 4), actin (10%, n= 4), and DNA (10%, n= 3) channels contributed the majority of population-specific effects, implicating intrinsic cell-type differences in cytoskeleton architecture, organelle morphology and metabolic state.

Thirty-six percent of DINO-merged PCs were associated with both timepoint and population, with the strongest effects in PC4 (R^2^_diff-timepoint_= 10%, p_adj_= 1×10^−37^ and R^2^_diff-population_= 7%, p_adj_= 1×10^−30^, **Fig. 3E-F**). Representative images indicated that PC4 captured cell and organelle polarization, which is also reflected in its association with CellProfiler feature sets describing cell and nuclear shape as well as the spatial organisation of the ER, mitochondria, tubulin and CD25 (**Suppl. Fig. 7E-F**). Polarization is a hallmark of T cell activation^48,49^, and temporal dynamics indicated a faster polarization in memory cells, consistent with their accelerated activation kinetics (**Fig. 3F**)^50,51^. The individual channel embeddings also uncovered organelle-specific changes associated with activation. For example, in the ER channel, PC2 captured increased ER complexity in memory cells during late activation (R^2^ = 9% with p_adj_= 7×10^−32^, R^2^ = 4% with p_adj_= 1×10^−7^, **Fig. 3G**).

Although DINO embeddings are not inherently interpretable, the recovery of these well-established biological patterns provides validation that they capture meaningful variation associated with trajectories of T cell activation and more subtle, cell-type-specific programs.

### Morphological clustering uncovers activation-dependent T cell states

To identify morphological and functional signatures of T cell activation, we applied Louvain clustering to the first 20 DINO-merged PCs, resolving nine clusters (**Fig. 3H**). Each cluster was annotated by CellProfiler features (**Fig. 3I, Suppl. Tab. 6**), cell cycle proportions (**Fig. 3J**) and representative images (**Fig. 3K**). Modelling naive and memory cell proportion per donor across activation revealed temporal dynamics (**Fig. 3L, Methods**). At rest, naive cells were enriched in Cluster 2 (93±1%), whereas memory cells were spread between Clusters 2 (33±9%) and 3 (50±4%). Cluster 3 differed from Cluster 2 by including a higher proportion of cycling cells (32% vs 7% in G1), having slightly higher expression of CD25, larger and less circular cells and more polarized ER and tubulin; Cluster 3 vs Cluster 2 β_cell_Intensity_MeanIntensity_CD25_= 0.6, β_cell_EquivalentDiameter_= 0.6, β_cell_FormFactor_= –1.2, β_ER_MassDisplacement_= 0.9, β_RadialDistribution_ZerkinkeMagnitude_2_0_tubulin_= 0.9 (all with p_adj_< 10×10^−324^, showcasing interpretable features of significant feature sets). This cluster expanded in early activation (12h) for both naive (34±6%) and memory cells (50±4%), before decreasing by 48h (naive: 16±4%, memory: 32±3%) and 120h (naive: 14±2%, memory: 16±4%), therefore representing a transient early-activation or primed state.

As activation progressed, the proportion of resting Cluster 2 decreased (timepoint β= –0.3, p= 6×10^−9^), giving rise to five activation-associated clusters (Clusters 1, 4, 6, 7 and 8) marked by canonical markers of activation: reduced frequency of G0 (< 12%), increased CD25 expression and, apart from Cluster 4, increased cell size: β_cell_intensity_mean_cd25_= 0.3-0.5 (p_adj_ < 7×10^−126^), β_cell_EquivalentDiameter_= 0.6-2.0 (p_adj_ < 9×10^−324^). Cluster 1 increased markedly over activation time (timepoint β= 0.2, p= 7×10^−6^), appearing earlier in memory (12h) than in naive cells (48h) (population:timepoint interaction β= 0.1, p= 3×10^−3^). It consisted mostly of cycling cells (76% G1, 20% S/G2/M) and was defined by irregularly-shaped large cells with high lipid and ER content and a large cytoskeleton: β_cell_AreaShape_FormFactor_= –0.5, β_cell_AreaShape_EquivalentDiameter_= 0.6, β_cyto_IntegratedIntensity_lipids_= 0.8, β_cyto_IntegratedIntensity_ER_= 0.7, β_cyto_IntegratedIntensity_tubulin_= 0.7 and β_cyto_IntegratedIntensity_actin_= 0.7 (all at p_adj_< 9×10^−324^, showcasing interpretable features of significant feature sets). At the further end of the activation trajectory, Clusters 7 and 8 represented the most activated states with the largest proportion of cells in S/G2/M phase (61% and 50%, respectively), and the largest, most irregular cells rich in mitochondria: β_cell_AreaShape_Area_= 1.4 and 2.0, β_cell_AreaShape_FormFactor_= –0.5 and –1.3 and β_cell_IntegratedIntensity_mito_= 1.3 and 0.9, respectively (all at p_adj_< 3×10^−163^, showcasing interpretable features of significant feature sets). These clusters were more frequent in memory than in naive cells peaking respectively at 48h (2±1% and 10±8%) and 120h (1±1 and 2±1%, respectively). Finally, Cluster 5 represents cells that are likely apoptotic, **w**ith small smooth-textured nuclei, and the lowest stain intensities observed in most stains **(Fig. 3I).** It was most abundant in naive cells at 120 and 168 hours after activation, but showed a slight increase in memories (**Fig. 3L, Suppl. Tab. 6).**

To further examine the morphologies that differentiate naive and memory cells throughout activation, we modelled all 465 CellProfiler feature sets for population effects and timepoint interactions (**Suppl. Tab. 7**). We identified 321 (69%) feature sets that either significantly differentiated the two populations at rest or showed different dynamics over time, indicating temporal divergence between naive and memory cells (**Fig. 3M-N)**. At baseline (0h), 225 feature sets distinguished memory from naive cells, with the most pronounced differences in mitochondria intensity and distribution, actin intensity and distribution, ER texture and distribution, cell or nucleus AreaShape and lipid intensity and granularity: β_mito_Intensity_1_= 1.5 (p_adj_= 4×10^−6^), β_mito_RadialDistribution_6_= 1.3 (p_adj_= 1×10^−10^), β_actin_RadialDistribution_6_= 1.3 (p_adj_= 5×10^−14^), β_actin_Intensity_1_= 0.8 (p_adj_= 4×10^−3^), β_ER_Texture_5_= 0.7 (p_adj_= 2×10^−11^), β_ER_RadialDistribution_13_= 0.7 (p_adj_= 10×10^−19^), β_cell_AreaShape_AreaShape_10_= 0.6 (p_adj_= 1×10^−19^), β_lipids_intensity_1_= 0.7 (p_adj_= 2×10^−3^) and β_lipids_Granularity_9_= –0.45 (p_adj_= 2×10^−5^) (**Fig. 3M**). Upon activation, the population differences diverged over time (**Fig. 3N**). In early activation, few feature sets (n= 74) showed a significant interaction term indicating that the baseline differences of naive and memory cells at rest were preserved after 12h of activation, except in the cell shape (n= 20), mitochondria intensity (n= 4) and actin radial distribution (n= 13) compartment. These patterns fit with population-specific activation speeds and cytoskeleton and mitochondria remodelling in early activation^24,52^. From 48h onwards, at which point the memory cells should have undergone at least one cell division, there were a number of feature sets across all channels that had a significant interaction: 193 (48h), 185 (120h) and 213 (168h). While memory T cells are known to respond more rapidly than naive cells to antigen stimulation, this analysis uncovers the complexity of the morphological landscape that reflects and shapes this readiness.

Together, TGlow resolves morphological and functional patterns associated with T cell activation and subtle population differences between naive and memory CD4^+^ T cells, highlighting the power of integrating deep learning approaches with classical feature extraction to uncover these phenotypes.

### CRISPR perturbations reveal gene-specific morphological signatures of T cell activation

A key challenge in high-throughput CRISPR screens is translating molecular perturbations to changes in cell phenotypes. We hypothesised that TGlow could bridge this gap by revealing whether gene perturbations generate distinct morphological and functional phenotypes or reposition T cells within a defined activation trajectory. To test this, memory CD4^+^ T cells from three healthy donors were CRISPR/Cas9-edited targeting four genes (*IL2RA, LCP2, CD69* and *LRRC32*) and an *AAVS1* cutting control (**Fig. 4A-B, Methods**). T cells were profiled by TGlow at baseline (0h) and after 16h of aCD2/CD3/CD28 re-stimulation (**Suppl. Fig. 8A**). Based on their function, knockout (KO) of *IL2RA* (encodes CD25 and is a key regulator of cell survival and proliferation) is expected to exert functional consequences at both resting and under stimulation, and KO of *LCP2* (encodes the adaptor protein SLP-76 mediating TCR/CD3 signalling required for cytoskeleton remodeling) is expected to manifest only upon activation. The KOs of *CD69* (encodes an adhesion molecule that is a marker of early activation) and *LRRC32* (encodes a protein that binds TGFb at the cell surface, primarily expressed in regulatory T cells) are expected to have minimal effects in this system and were used as negative controls to gauge false positive rate due to technical effects.

**Figure 4.**
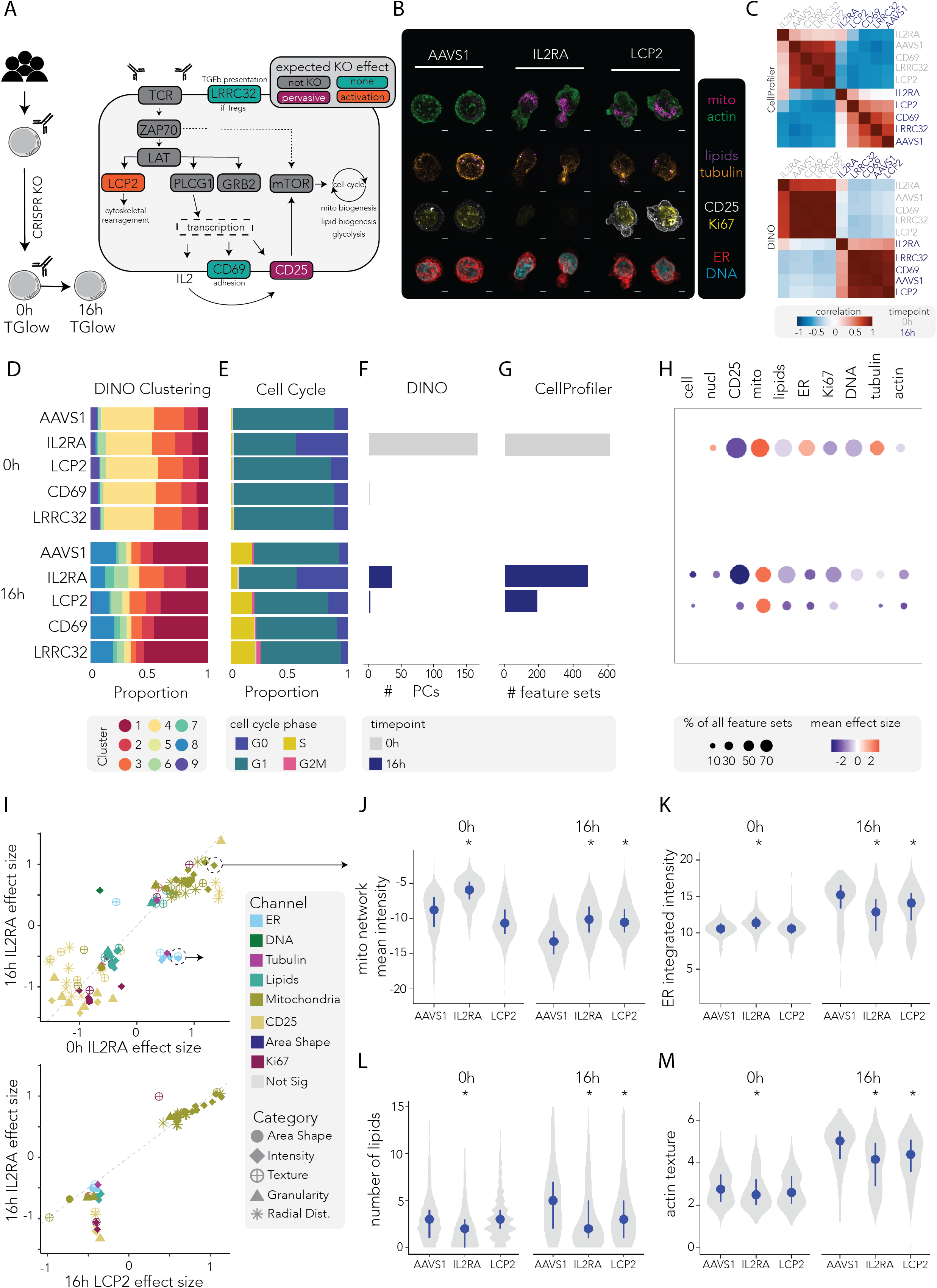
CRISPR knockouts of T cell activation genes reveal gene-specific morphological signatures captured by TGlow. **A)** Left: experimental design; Right: schematic of relevant signalling pathways of T cell activation, with the targeted genes colored by their expected phenotype: no expected phenotype (teal: LRRC32, CD69), pervasive phenotype at 0h and 16h (burgundy: IL2RA), phenotype upon activation (orange: LCP2) and not targeted (grey). **B)** Representative images (cell closest to the centroid of the sample and its closest neighbor, deconvoluted and max projected) of 16h cells for *AAVS1*, *IL2RA* and *LCP2* (n= 2 each, columns) colored by four combinations (rows): mitochondria (pink) and actin (green); lipids (purple) and tubulin (orange); CD25 (white) and Ki67 (yellow) or ER (red) and DNA (blue). Scale bar= 2.5μm. **C)** Heatmap of sample correlation with transformed and scaled CellProfiler features (top) or unscaled DINO-PCs (bottom). Features or PCs are averaged by timepoint_target combination. Timepoints: 0h (text in grey) and 16h (text in dark blue). Correlation: –1 (blue) to 0 (white) to +1 (red). **D-H)** Plots per target (y-axis) and timepoint (0h: top panel, 16h: bottom panel) showing **D)** the proportion of cells in each cluster of Figure 3H) (n= 9) when edited cells are jointly clustered with the reference timecourse; **E**) the proportion of cells in each cell cycle phase: G0 (blue), G1 (green), S (yellow) or G2M (pink); **F)** the number of significant differential DINO-PCs, colored by timepoint (0h: grey, 16h: blue), **G)** the number of significant differential CellProfiler feature sets, colored by timepoint (0h: grey, 16h: blue) and **H)** feature plot separated by target/timepoint combination (row) and channel (n= 9, column). The size of the dot represents the percentage of significant feature sets in that channel and the color depicts the mean effect size of the significant feature sets: –2 (blue) to 0 (white) and +2 (red). **I)** Dotplot showing the correlation between the mean effect sizes of significant feature sets shared between *IL2RA*-KO at 0h and 16h (top) or *IL2RA*-KO and *LCP2*-KO at 16h (bottom). The dots are colored by channel (ER: light blue, DNA: green, tubulin: purple, lipids: teal, mitochondria: green, CD25: yellow, AreaShape: dark blue, Ki67: burgundy) and shaped by category (AreaShape: large circle, Intensity: diamond, Texture: circle and cross, Granularity: triangle, Radial Distribution (Radial Dist.): star). **J-M)** Single-cell violin plots of the transformed and scaled features with the median and interquantile range for each perturbation (*AAVS1*, *IL2RA*, *LCP2*) at each timepoint (0h: left, 16h:right). Significant differences (p_adj-LRT_ ≤ 0.05) are shown with an asterix (*). The features are: **I)** mitoNetwork_Intensity_MeanIntensity, **K)** cell_Intentisy_IntegratedIntensity_ER, **L)** cell_Children_lipids_Count and **M)** cell_Texture_Contrast_actin_3_00_256.

First, pairwise correlations of imaging profiles across timepoint–gene target combinations were computed using either DINO-PCs or CellProfiler features (**Fig. 4C, Suppl. Fig. 8F-G**). For both analyses, samples clustered strongly by timepoint, indicating that activation status is the dominant source of variation in the data. Within each timepoint, *IL2RA* knockout consistently exhibited the lowest correlation with other perturbations, reflecting its pronounced effect on T cell states. Notably, the *IL2RA* knockout at 16h maintained a high similarity to *IL2RA* knockout at 0h, suggesting a partial failure to transition into a fully activated phenotype upon stimulation. This observation is consistent with the expected role of IL-2 signaling in sustaining activation-induced remodeling. Within the CellProfiler feature space, *LCP2* knockout profiles showed greater similarity to *IL2RA* knockouts than to other perturbations, specifically under stimulated conditions. This suggests partial convergence toward a low-activation phenotype, consistent with *LCP2* acting through a distinct, stimulation-dependent mechanism. Such analyses confirm that TGlow captures gene-specific morphological signatures rather than generic perturbation responses.

Joint clustering of the CRISPR-edited cells with the reference activation trajectory (**Fig. 3H**, **Suppl. Fig. 8B**) recapitulated expected activation dynamics. At 0h, *AAVS1*-edited cells were distributed among resting Clusters 2 and 3 (mean= 11±1% and 24±2%, respectively) and activated Clusters 4, 6 and 1 (mean= 47±5%, 3±1%, 8±0%, respectively), consistent with the prior *in vitro* activation history of these cells (**Fig. 4D**). Upon restimulation, *AAVS1*-edited cells shifted towards the activated Clusters 1, 6, 7 and 8 (mean= 46±1%, 7±5%, 3±1% and 20±3%, respectively). By contrast, *IL2RA*-deficient cells failed to follow this activation trajectory, with reduced representation of the activated Clusters 1, 6, 7 and 8 (mean= 17±7%, 13±2, 7±3 and 11±7%, respectively), and expansion of the resting Clusters 2, 3 and 4 (mean= 22±16%, 21±3% and 9±4%, respectively). This impaired activation was corroborated by cell cycle arrest, with increased cells in G0 at both 0h (44±16% *IL2RA* vs 11±2% *AAVS1,* β= 33%, p= 0.02) and 16h (48±23% *IL2RA* vs 7.5±1% *AAVS1*, β= 41%, p= 0.04) (**Fig. 4E, Suppl. Fig. 8C-D**), consistent with the role of *IL2RA* in promoting survival and proliferation^53^.

To identify gene-specific morphological changes, we fitted linear mixed models comparing each perturbation to *AAVS1* at each timepoint using either DINO PCs (**Fig. 4F, Suppl. Tab. 8**) or CellProfiler features (**Fig. 4G-H, Suppl. Tab. 9**). As expected, *CD69* and *LRRC32* KO showed minimal effects, showing no differential CellProfiler features and only one differential DINO PC for *CD69* at 0h (lipids-PC31 β= 0.1, p= 0.04, R^2^ = 0.07%). By contrast, *IL2RA* KO induced pervasive morphological changes across timepoints affecting 196 (40%) and 134 (27%) CellProfiler feature sets and 166 (26%) and 35 (5%) DINO PCs at 0h and 16h, respectively. The perturbed features spanned all imaging channels with the strongest effects in the CD25 and mitochondria compartments, reflecting mitochondrial dysfunction reported in *IL2RA*-deficient regulatory T cells^54^: CD25-Intensity feature sets: 100% impacted at 0h (mean β= –0.9, p_adj_= 2×10^−7^) and 67% at 16h (mean β= – 1.2, p_adj_= 2×10^−3^) and mito-Intensity: 77% impacted at 0h (mean β= 0.9, p_adj_= 7×10^−3^) and 61% at 16h (mean β= 0.8, p_adj_= 1×10^−2^, **Fig. 4H-J**).

Timepoint-dependent effects were evident in several compartments, particularly in the lipids, actin, ER and AreaShape: lipid-AreaShape feature sets: 62% affected at 0h (mean β= –0.2, p_adj_≤ 2×10^−3^) and 12% at 16h (mean β= 0.6, p_adj_≤ 2×10^−4^); actin-RadialDistribution feature sets: 18% affected at 0h (mean β= –0.1, p_adj_≤ 3×10^−2^) and 11% at 16h (mean β= –0.4, p_adj_≤ 0.03); actin-Texture feature sets: 20% affected at 0h (mean β= –0.005, p_adj_≤ 5×10^−2^) and 40% at 16h (mean β= 0.4, p_adj_≤ 0.03); tubulin-Intensity feature sets: 60% affected at 0h (mean β= 0.6, p_adj_≤ 8×10^−3^) and 20% at 16h (mean β= 0.1, p_adj_≤ 0.03); tubulin-RadialDistribution feature sets: 22% affected at 0h (mean β= 0.4, p_adj_≤ 1.4×10^−2^) and 6% at 16h (mean β= –0.009, p_adj_≤ 0.04); tubulin-Texture: 38% affected at 0h (mean β= –0.2, p_adj_≤ 1×10^−3^) and 0% at 16h; ER-Intensity feature sets: 80% affected at 0h (mean β= 0.5, p_adj_≤ 1×10^−2^) and 33% at 16h (mean β= –0.5, p_adj_≤ 0.01); ER-RadialDistribution feature sets: 39% affected at 0h (mean β= 0.3, p_adj_≤ 3×10^−3^) and 0% at 16h; ER-Granularity feature sets: 0% affected at 0h and 27% at 16h (mean β= –0.4, p_adj_≤ 0.02); and nucleus-Areashape feature sets: 5% affected at 0h (mean β= –0.4, p_adj_≤ 6×10^−3^) and 7% at 16h (mean β= 0.3, p_adj_≤ 0.01, **Fig. 4I, top panel**). The most pronounced timepoint-dependent effect occurred in the ER, where *IL2RA*-deficient cells showed greater ER abundance at rest, but failed to upregulate it following stimulation (**Fig. 4K**), suggesting ER stress or metabolic adaptation. Dysregulated lipid biogenesis further supported altered metabolic adaptation in *IL2RA*-deficient cells (**Fig. 4L**), echoing CD25-dependent lipid biosynthesis in regulatory T cells^54^. These findings demonstrate that *IL2RA*-deficiency disrupts key metabolic programs involving the mitochondria, ER and lipid compartment, ultimately impairing T cell survival and function.

Consistent with its role as a TCR-proximal adaptor, *LCP2* KO induced morphological perturbations only after restimulation, with 52 (11%) differential CellProfiler feature sets and 2 DINO PCs detected exclusively at 16h. As expected due to its role in controlling cytoskeleton rearrangement, activated *LCP2*-deficient cells exhibited disrupted actin and tubulin organisation: impacted were 33% of actin-Intensity feature sets (mean β= –0.5, p_adj_= 0.04), 4% of actin-RadialDistribution feature sets (mean β= –0.4, p_adj_= 0.03) and 13% of tubulin-Intensity feature sets (mean β= –0.4, p_adj_= 0.03, **Fig. 4M**)^55–57^.

Though we did not detect any significant effect on cluster or cell cycle composition (**Suppl. Fig. 7D**), *LCP2*-deficient cells exhibited other hallmarks of impaired T cell activation such as decreased expression of CD25 and Ki67 expression, with effect sizes smaller than those observed in *IL2RA*-deficient cells: mean β_CD25_Texture_= –0.4 (p_adj_= 0.05), mean β_CD25_Intensity_= –0.4 (p_adj_= 0.05), mean β_KI67_Intensity_= –0.4 (p_adj_= 0.04), (**Fig. 4I**, bottom panel)^58,59^. Other organelles mirrored the effects of *IL2RA* KO, such as increased mitochondria activity, reduced ER and altered lipids: mean β_mito_Intensity_= 0.9 (p_adj_= 0.03), mean β_mito_RadialDistribution_= 0.6 (p_adj_= 0.02), mean β_mito_AreaShape_= 0.3 (p_adj_= 0.02), β_mito_Texture_= 0.2 (p_adj_= 0.01), β_ER_Intensity_= –0.4 (p_adj_= 0.03), β_ER_Texture_= –0.4 (p_adj_= 0.04), β_ER_Granularity_= –0.4 (p_adj_= 0.03), β_lipids_Intensity_= –0.4 (p_adj_= 0.03) and β_lipids_AreaShape_= –0.4 (p_adj_= 0.03). These results indicate that *LCP2* disruption impairs activation through cytoskeletal and metabolic pathways that are partly shared with, but mechanistically distinct from, those regulated by *IL2RA*.

We thus establish TGlow as a sensitive and scalable platform for mechanistic phenotyping of genetic perturbations. The mapping of CRISPR-edited cells into the activation reference further illustrates that reference maps derived from TGlow can be extended across datasets, enabling comparative phenotyping at scale.

### TGlow resolves morphological hallmarks of CD8^+^ T cell exhaustion

TGlow’s high-content, multidimensional view of cellular architecture, metabolism and function enables it to be easily repurposed beyond CD4^+^ T cell activation to profile diverse immune cell states. To test this capacity, we applied TGlow to a model of CD8^+^ T cell exhaustion, a dysfunctional state arising from chronic antigen exposure marked by loss of effector function, metabolic rewiring and proliferative arrest. To model exhaustion, CD8^+^ T cells were activated and either expanded for eight days or repeatedly restimulated every 48 hours to mimic persistent antigen exposure (**Fig. 5A**)^60^. TGlow yielded 484 CellProfiler feature sets for 6,239 expanded and 3,317 exhausted CD8^+^ T cells (**Fig. 5B-C, Suppl. Tab. 2**).

**Figure 5.**
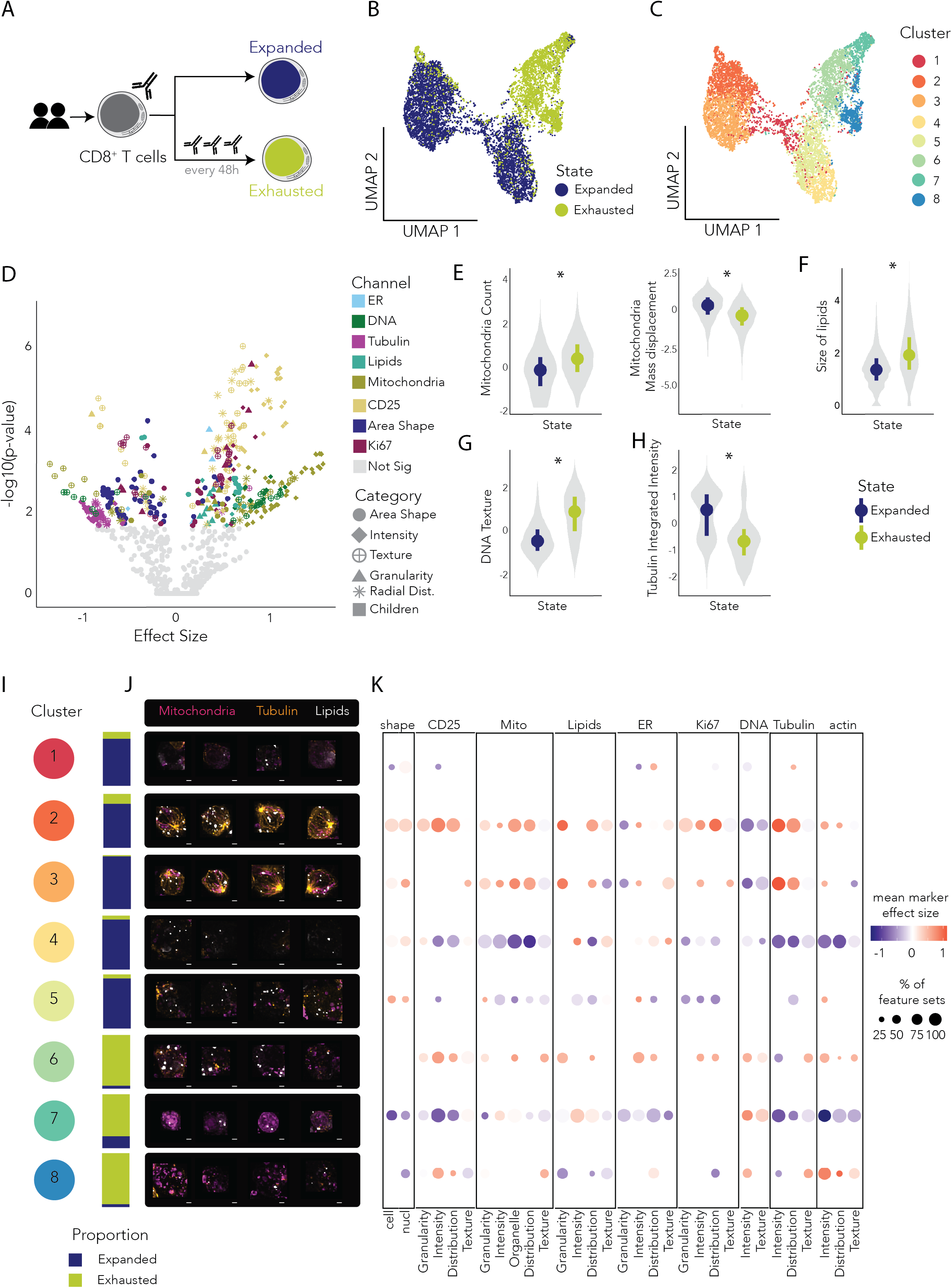
TGlow uncovers known and unknown morphologies of CD8^+^ T cell exhaustion. **A)** Experimental design. **B-C)** UMAP embeddings of 9,556 cells (15 PCs) which are annotated by: **B)** state (expanded: blue, exhausted: green) or by **C)** Louvain cluster number (1-8, resolution= 0.8). **D)** Volcano plot displaying the effect size (x-axis) versus the –log10(p-adjusted) from modelling CD8^+^ T cell states with expanded cells as a reference. Each dot represents the mean of a feature set, with grey circles indicating no statistical dysregulation and otherwise, colors indicating channels (ER: light blue, DNA: green, tubulin: purple, lipids: teal, mitochondria: green, CD25: yellow, AreaShape: dark blue, Ki67: burgundy) and shapes indicating categories (AreaShape: large circle, Intensity: diamond, Texture: circle and cross, Granularity: triangle, Radial Distribution (Radial Dist.): star). **E-H)** Single-cell violin plots of the transformed and scaled features with the median and interquantile range for each status indicated in a coloured point and line: expanded (blue) and exhausted (green). Significant differences (p_adj_ ≤ 0.05) are shown with an asterisk (*). The features are: **E)** cell_Children_mito_Count (mitochondria count, left) and cell_Intensity_MassDisplacement_mito (mitochondria mass displacement, right), **F)** lipids_AreaShape_Area (size of lipids), **G)** nucl_Texture_Variance_dna_3_00_256 (DNA texture) and **H)** cell_Intensity_IntegratedIntensity_tubulin (tubulin integrated intensity). **I-K)** Connected plots summarising each cluster (row): **I)** proportion plot of expanded (blue) and exhausted (green) cells **J)** Representative images (n= 4, cell closest to the cluster centroid and its three nearest neighbours, deconvoluted and max projected) showing mitochondria (pink), tubulin (orange) and lipids (white). Scale bar= 2.5μm. **K)** Marker plot where the size of the dot represents the percentage of significant feature sets per cluster (row) in each channel (column) and the color depicts the mean marker effect size from –1 (blue) to 0 (white) and +1 (red).

Linear mixed modelling identified 169 feature sets (35%) that strongly discriminated between expanded and exhausted CD8^+^ T cells, encompassing both canonical hallmarks of exhaustion and previously unrecognized phenotypes (**Fig. 5D, Suppl. Tab. 10**). The perturbations spanned multiple cellular processes involving mitochondria, DNA, lipids, CD25, Ki67, cell or nucleus AreaShape and tubulin: mean β_mitochondria_= 0.7 (54% of mitochondria feature sets), mean β_DNA_= 0.5 (17%), mean β_lipids_= 0.4 (27%), mean β_CD25_= 0.4 (77%), β_Ki67_= 0.2 (62%), mean β_AreaShape_= –0.4 (44%) and mean β_tubulin_= –0.8 (54%). More specifically, and consistent with prior reports, exhausted CD8^+^ T cells contained more mitochondria that failed to polarize effectively: β_cell_Children_mito_Count_= 0.5 (p_adj=_ 0.008) and β_cell_Intensity_MassDisplacement_mito_= –0.7 (p_adj=_ 0.003) (**Fig. 5E**)^60–63^. Furthermore, exhausted T cells contained fewer, but larger lipid droplets: β_cell_Children_lipids_Count_= –0.5 (p_adj_= 0.04) and β_lipids_AreaShape_Area_= 0.7 (p_adj_= 2 × 10^−3^, **Fig. 5F**)^64–66^. This is corroborated by exhausted cells showing higher values for high granularity measurements (2 to 8), consistent with overall larger lipid structures (**Suppl. Fig. 9A**).

They also displayed a higher DNA content and texture, consistent with large-scale chromatin reorganization^67^: 66% of DNA_intensity features sets (mean β= 0.9, p_adj_≤ 0.01) and 86% of DNA_texture sets (mean β= 0.3, p_adj_≤ 0.01) with *nucl_Texture_DifferenceEntropy_dna_3_00_256* (β= 1.1, p_adj_= 3×10^−5^) as an illustrative feature of the significant feature set *DNA_texture 2* (**Fig. 5G**).

Strikingly, we observed a near-complete loss of tubulin architecture with disruptions across the tubulin_intensity, _texture and _RadialDistribution feature sets. Respectively, 83% (mean β= –0.9, p_adj_≤ 0.02), 67% (mean β= –0.7, p_adj_≤ 0.01) and 33% (mean β= –0.7, p_adj_≤ 0.02) of feature sets were impacted. This response is exemplified by *cell_Intensity_IntegratedIntensity_tubulin* (β= –1.1, p_adj_= 0.02), an interpretable feature of the set *tubulin_intensity 2* (**Fig. 5H**) and was coordinated with mitochondria repositioning (**Suppl. Fig. 9B**). This cytoskeletal phenotype has not been previously described in T cell exhaustion.

We further explored CD8^+^ T cell heterogeneity within the expanded and exhausted populations, identifying eight clusters with distinct morphological features (**Fig. 5C,I-K, Suppl. Tab. 11**). Five clusters were predominantly composed of expanded cells: Clusters 1(87%), 2 (81%), 3 (98%), 4 (90%) and 5 (92%) (**Fig. 5I**). In the expanded population, Cluster 2 featured proliferating cells with high Ki67 expression and Cluster 1 displayed features typical of dying cells: Cluster 2 β_nucl_Intensity_IntegratedIntensity_ki67_= 1.3 (p_adj_ < 10×10^−324^); Cluster 3 β_cell_AreaShape_MeanRadius_= –0.7 (p_adj_= 3×10^−69^), β_nucl_AreaShape_FormFactor_= –0.8 (p_adj_= 2×10^−100^) and β_nucl_Texture_Correlation_dna_3_00_256_= 0.6 (p_adj_= 8×10^−60^). The remaining expanded cells separated in either Cluster 3 with abundant tubulin, numerous lipid droplets and a large polarized mitochondria network or clusters with less abundant tubulin (Clusters 4 and 5): Cluster 3: β_cell_Intensity_IntegratedIntensity_tubulin_= 1.4 (p_adj_ < 10×10^−324^), β_cell_Children_lipids_Count_= 0.9 (p_adj_= 3×10^−300^), β_mitoNetwork_AreaShape_EquivalentDiameter_= 1.0 (p_adj_= 3×10^−162^) and β_cell_Intensity_MassDisplacement_mito=_ 0.8 (p_adj_= 7×10^−192^). Whilst Cluster 5 represented the largest cells in the data and displayed higher mitochondria granularity, Cluster 4 had fewer mitochondria and a smoother actin cytoskeleton: Cluster 5: β_cell_AreaShape_EquivalentDiameter_= 0.6 (p_adj_= 3×10^−11^) and β_cell_Granularity_1_mito_= 0.8 (p_adj_= 9×10^−126^); Cluster 4: β_cell_Children_mito_Count_= –1.4 (p_adj_= 2×10^−93^), β_cell_Texture_InverseDifferenceMoment_actin_3_00_256_= 1.2 (p_adj_= 3×10^−314^) and β_cell_Intensity_MeanIntensity_actin=_ –0.9 (p_adj_= 2×10^−159^).

Three clusters were predominantly composed of exhausted cells: Clusters 6 (95%), 7 (79%) and 8 (96%) (**Fig. 5I**). Among the exhausted cells, Clusters 6 and 8 displayed the highest tubulin content, with levels similar to that of expanded Cluster 4 (the lowest tubulin content in the expanded cluster): β_cell_Intensity_MeanIntensity_tubulin_= –0.4 (p_adj_= 2×10^−110^) and –0.3 (p_adj_= 10×10^−30^), respectively. The other top features marking Cluster 6 were high expression of CD25 and ER content, with moderate Ki67 expression, possibly indicating a proliferating cluster of exhausted cells: β_memb_Intensity_IntegratedIntensity_cd25_= 0.8 (p_adj_= 9×10^−169^), (β_cyto_Intensity_IntegratedIntensity_ER_= 0.7 (padj= 2×10^−117^) and β_nucl_Intensity_MeanIntensity_ki67_= 0.5 (p_adj_= 4×10^−45^). The cells in Cluster 8 were high in actin, CD25 expression and had the fewest lipid droplets in the dataset suggesting reprogramming of membrane and metabolic architecture: β_cell_Intensity_MeanIntensity_actin_= 1 (p_adj_= 7×10^−108^), β_memb_Intensity_IntegratedIntensity_cd25_= 0.8 (p_adj_= 2×10^−86^) and β_cell_Children_lipids_Count_= –0.6, p_adj_= 5×10^−39^). Finally Cluster 7 was the most depleted of tubulin and grouped the smallest cells in the dataset, but most metabolically active exhausted cells cells with high DNA content, possibly marking dying exhausted CD8^+^ T cells: β_cell_Intensity_MeanIntensity_tubulin_= –0.7 (p_adj_= 2×10^−240^), β_cell_AreaShape_EquivalentDiameter_= –1.9 (p_adj_< 10×10^−324^), β_cell_Intensity_MeanIntensity_mito_= 0.9 (p_adj_= 1×10^−252^) and β_nucl_Intensity_MeanIntensity_dna_= 0.7 (p_adj_= 4×10^−76^).

Hence, TGlow extends beyond CD4^+^ T cell activation by not only recapitulating hallmarks of impaired metabolism in CD8^+^ T cell exhaustion but also highlighting novel insights into impaired cytoskeleton architecture associated with CD8^+^ T cell exhaustion. TGlow is a broadly applicable platform for quantitative mapping of morphological and functional states across lymphocyte lineages.

## Discussion

TGlow is an end-to-end image-based profiling ecosystem for quantifying lymphocytic phenotypes with single-cell resolution and at scale. It unites an optimized experimental protocol with an open-source computational workflow enabling high-throughput sample preparation, image processing, feature extraction, and data analysis. Although developed for T cells, TGlow is readily adaptable to other cell types, marker panels, and analytical strategies.

In contrast to traditional, hypothesis-driven and low-throughput lymphocyte imaging, TGlow, like other HCI approaches, treats microscopy as a discovery platform for systematic and unbiased phenotypic profiling. While a few studies have applied assays such as Cell Painting to lymphocytes, none, to our knowledge, have optimized for lymphocyte-specific biological questions or the technical challenges they present while maintaining resolution, scalability and data richness^68–71^. TGlow’s dense z-sampling captures the size diversity of lymphocytes (5–15 μm), critical for mapping heterogeneous lymphocyte states, and improves optical resolution through deconvolution. Building upon the success of cell-type specific adaptations of CellPainting^6,72,73^, TGlow incorporates cyclic imaging to profile a broader range of lymphocyte markers. This design renders TGlow uniquely capable of jointly profiling cellular and organelle morphology, metabolism, and function to provide a more extensive view of coordinated lymphocyte behavior.

TGlow faithfully recovers hallmark morphological, metabolic and functional states associated with cell-type identity, activation, exhaustion, and drug or genetic perturbations. The discrimination of naive and memory CD4^+^ T cells, which differ in their immunological history and capacity for effector functions, exemplifies this: TGlow captures enriched mitochondrial content, reorganized cytoskeleton architectures, and compartment complexity in memory cells that represent structural and functional features likely underpinning their rapid mobilization and multi-effector capacity^76–79^. These results reinforce a view of morphology, metabolism and function as both a readout and regulator of cellular state^40,70,80^. More broadly, TGlow’s capacity to resolve such subtle but functionally meaningful states opens new opportunities to link heterogeneous or dysfunctional morphological and metabolic states to function across immune contexts.

Despite its advantages, TGlow has practical and analytical constraints. High-content confocal imaging requires specialized microscopy instrumentation, substantial data storage, and computational resources that may limit accessibility. Lymphocyte adherence remains challenging, with cell losses (5-75%) between imaging cycles largely driven by delays in staining and acquisition, which in practice cannot always be avoided due to equipment use or failures. Batch effects at the well or plate level, arising from fluctuations in reagent performance, acquisition parameters or illumination, present further reproducibility and scalability challenges. Computational harmonization tools such as Harmony or Combat^81–83^ can mitigate imaging-derived batch effects, but typically assume comparable samples, rendering them unsuitable for time-resolved datasets where batch is co-linear with time. To reduce batch variability, TGlow scales image intensities to plate controls (Jurkats) and includes random effect terms to model well level variation, though biological drifts and variation from prolonged cell culture may limit their use as a stable reference. Engineered, fixed or drug-treated cells could offer a more robust normalization control.

The interpretability of both classical and deep-learning derived imaging features remains an open challenge. While some metrics correlate with known processes, such as abundance, size or polarization, many capture abstract representations of high-order spatial relationships that are difficult to interpret. Mapping these signatures to biological meaning will require dedicated validation through perturbation experiments and integration with orthogonal modalities such as transcriptomics, proteomics, and functional assays. Further improvements could be gained by extending TGlow’s feature extraction pipeline toward 3D analysis, resolving phenotypes that are currently masked by max-projection, such as include nuclear invaginations, organelle colocalization (i.e: ER-mitochondria) or network structure that provide additional mechanistic insights into lymphocyte biology^70,84^. Although TGlow’s pipeline supports 3D analysis, the computational burden of 3D feature extraction in CellProfiler makes it currently prohibitive to do at large scales. Finally, because TGlow is optimized for suspension cells, it cannot yet capture the spatial organization and cell-cell interactions intrinsic to tissue environments. Expanding its use in co-cultures or tissues would more fully represent lymphocyte behavior *in vivo*.

TGlow’s core strength lies in its ability to quantify unbiased and biologically relevant phenotypes of lymphocytes at scale, positioning it as a powerful platform for discovery-driven immunological research. It is especially suited for large perturbation screens and population-scale profiling, where imaging can not only serve as a cost-effective complement to single-cell omics, but also bridge molecular signatures with cell behavior. Integration of TGlow with multi-omic modalities will help define the bidirectional relationship between molecular programs and cellular phenotypes, enabling more accurate predictive modeling of immune cell behavior. This positions TGlow as a key component of genetic efforts to map variants-to-function, profiling how genetic variation shapes lymphocyte morphology and function beyond molecular regulation. In summary, TGlow broadens high-content imaging for lymphocytes through its unique combination of scalability, resolution and multiplexing to deliver morphological and functional insights at single cell resolution, paving the way for the next generation of immune profiling.

## Methods

### Cell culture

All cell cultures were kept at 37°C and 5% CO_2_ atmosphere. Cell counts were obtained on a Cellometer™ Spectrum image cytometer (Revvity) or the Cellaca^™^ MX high-throughput cell counter (Revvity) with AO/PI staining (Nexcelom CS2-0106).

### Cell lines

The Jurkat, Clone E6-1 cell line was purchased from ATCC (#TIB-152^TM^) and cultured in RPMI-1640 (Gibco^TM^, #11875093), supplemented with 10% heat-inactivated fetal bovine serum (FBS, Sigma-Aldrich^®^ #F9665-500ML), 2 mM L-glutamine (Gibco^TM^, #25030081), 1mM sodium pyruvate (Gibco^TM^, #11360070), 1X MEM non-essential amino acids (Gibco^TM^, #11140050), 100 U/ml penicillin and 100 µg/ml streptomycin (Gibco^TM^, #15140122). The density was maintained at 0.25 million cells per mL by splitting the cells 1:2 every two or three days. For cryopreservation, cell pellets containing 1 million cells were resuspended in 1 mL of complete growth medium supplemented with 10% DMSO. The cell suspensions were placed in Mr. Frosty (Nalgene) or CoolCell (Corning) overnight at –80°C and then transferred to liquid nitrogen for long-term storage. Cryo-preserved cells were thawed in a 37°C water bath and washed in 10mL of growth medium before cell culture.

### Human samples

Human samples were obtained ethically and their research use was in accordance with the terms of an institutional review board/ethics committee-approved protocol (15/NW/0282). Normal Human Leukopaks were purchased from BioIVT^®^, with inclusion criteria of gender female, age 20–40 years, BMI < 28, white caucasian ancestry and non-smoking status. The donors were distributed across the projects like so: Donor 1 (34 years old, BMI= 21.1 – Drug), Donor 2 (43 years old, BMI= 23.8; Drug + CRISPR), Donor 3 (21 years old, BMI= 26.4; Drug), Donor 4 (44 years old, BMI= 23.5; Timecourse + CRISPR), Donor 5 (36 years old, BMI= 23.7; Timecourse + CRISPR), Donor 6 (30 years old, BMI= 20.8; Timecourse), Donor 7 (29 years old, BMI= 24.5; Timecourse), Donor 8 (34years old, BMI= 27.4; Exhaustion) and Donor 9 (18years old, BMI= 28.1; Exhaustion).

### Peripheral blood mononuclear cells (PBMCs) isolation

Leukopaks were diluted 1:1 with 1 mM EDTA-1% FBS-RPMI, and PBMCs were isolated using density gradient centrifugation with Ficoll-Paque PLUS (Cytiva, #17-1440-02). A 25mL aliquot of the diluted sample was layered onto 17 mL of Ficoll-Paque Plus (Cytiva) in a 50mL Falcon tube and centrifuged at 800 x*g* for 30 minutes at room temperature. PBMCs were collected from the intermediate layer, washed four times with cold EasySep buffer (Stemcell Technologies, #20144) and cryopreserved at 100 million cells per mL in FBS with 20% DMSO. Cells were cooled overnight to –80°C in a Mr. Frosty or a CoolCell and then transferred to liquid nitrogen for long-term storage.

### CD4^+^ T cell isolation and cell culture

Cryo-preserved PBMCs were thawed in a 37°C water bath, washed and rested overnight at 20×10^6^ cells/mL in RPMI-1640 supplemented with 10% FBS, 2 mM L-glutamine, 100 U/ml penicillin and 100 µg/ml streptomycin. Naive and memory CD4^+^ T cells were isolated with the EasySep^TM^ Human Naive CD4^+^ T cell Isolation Kit II (#17555, STEMCELL Technologies^TM^) or the EasySep^TM^ memory CD4^+^ T cell enrichment kit (#19157, STEMCELL Technologies^TM^), following the manufacturer’s instructions. Isolated T cells were cultured in round-bottom 96-well plates or flat-bottom 24-well plates at 1×10^6^ cells/mL in StemPro^TM^34-SFM (Gibco^TM^, #10639011) supplemented with 10% FBS, 100 U/ml penicillin and 100 µg/ml streptomycin. Unless specified, recombinant human IL-2 (R&D Systems, #10453-IL) was supplemented every two days (15 ng/mL, 140U/mL). T cells were activated using ImmunoCult™ Human aCD3/CD28/CD2 T Cell Activator (STEMCELL Technologies^TM^, #10970) at 12µL per million cells.

### Drug perturbations

Memory CD4^+^ T cells were activated for four hours before treatment with the compounds listed in **Table 1**. The drug dilutions were prepared in a deep-well 96-well plate and then stamped with the ViaFlow 96 (Integra Biosciences AG) into 96-well cell culture plates before addition of the cells. Due to its reversible mechanism of action, palbociclib was added again at 28 and 52 hours post-activation. Vehicle controls (DMSO or H_2_O) and media-only controls (cStemPro medium only) were included for each donor in each cell culture and imaging plate. The samples were collected at 0, 24 and 72 hours post-activation for TGlow profiling.

**Table 1.** Compounds selection for the drug perturbation experiment. The table lists all compounds used, their suppliers and catalog numbers, solvents, and the range of concentrations tested. Each compound’s known mechanism of action is briefly summarized based on published literature. Mechanistic descriptions highlight key molecular targets and biological effects relevant to cell cycle control, mitochondrial function, or immune regulation

| Compound | Supplier | Catalogue # | Solvent | Concentrations (μM) |
| --- | --- | --- | --- | --- |
| Palbociclib | MilliporeSigma | PZ0383 | DMSO | 0.002, 0.02, 0.2, 1, 2 |
| Hydroxyurea (HU) | Sigma Aldrich | H8627 | H<sub>2</sub>O | 10, 100, 500, 1000, 5000 |
| Mycophenolic acid (MPA) | Sigma Aldrich | 89287 | DMSO | 0.03, 0.1, 0.3, 1, 3 |
| Disulfiram | Selleck Chemicals | S1680 | DMSO | 0.1, 0.5, 1, 2.5, 10 |
| Rotenone | Sigma Aldrich | 45656 | DMSO | 0.01, 0.1, 1, 5, 20 |
| mdivi-1 | Sigma Aldrich | M0199 | DMSO | 0.1, 1, 10, 50, 100 |
| MFI8 | MedChem Express LLC | HY-150031 | DMSO | 0.1, 1, 5, 20, 100 |
| Tofacitinib | MedChem Express LLC | HY-40354 | DMSO | 0.05, 0.1, 0.5, 1, 10 |

### T cell activation timecourse

Naive and memory CD4^+^ T cells were isolated as previously described and each population was activated as a pool before stamping with the ViaFlow 96 into five 96-well cell culture plates and cultured without IL-2. Timepoints were harvested at: 0 hours, 12 hours, 48 hours, 120 hours and 168 hours for TGlow profiling.

### CRISPR

Lyophilized sgRNAs (EditCo, 3 sgRNAs per target, (**Suppl. Tab. 12**) were resuspended in water to a stock concentration of 100µM and stored at –20°C until use. Ribonucleoproteins (RNP) were produced by mixing sgRNAs (180pmol per reaction) and Cas9 (Alt-R™ S.p. Cas9 Nuclease V3, IDT, 49pmol per reaction) in a v-bottom 96-well plate. Memory CD4^+^ T cells, activated for 48 hours (24µL of Immunocult per million cells), were collected and spun down for 5min at 400g then washed twice with DPBS without calcium or magnesium, before being resuspended in P3 Primary Cell 4D-Nucleofector® buffer (Lonza, cat. #V4XP-3032) at 3×10^5^ cells per 20µL (per reaction) and added to the RNP mix. The cells in the buffer were then transferred to a 16-well Nucleocuvette™ Strip (Lonza, 4D-Nucleofector™ X Kit) for nucleofection using the pulse code EH-115. Immediately after nucleofection, 80µL of pre-warmed cStemPro media supplemented with IL-2 (15 ng/mL, 140U/mL) was added to each well and incubated at 37°C with 5% CO2 for 30min. The cells were then transferred to a 96-well round-bottom plate containing 225µL of cStemPro supplemented with IL-2 (15 ng/mL, 140U/mL) and incubated at 37°C with 5% CO2 for 5 days. Edited cells were re-activated (12µL of Immunocult per million cells) and sampled at 0h and 16h after re-stimulation for TGlow profiling. Editing efficiencies were confirmed by Sanger sequencing (Genewiz^®^) and ICE analysis from EditCo^©^ (**Suppl. Tab. 12**).

### In vitro exhaustion protocol

CD8^+^ T cells were isolated from healthy donors using the REAlease CD8 MicroBead Kit (Miltenyi Biotec, #130-117-036) according to the manufacturer’s instructions. Cells were counted and seeded at a density of 1×10^6^ per mL in complete media: Tex MACS media supplemented with 1% human serum and 100 U/mL human IL-2 IS (Miltenyi Biotec, 130-097-748). Initial activation is with 1:100 T Cell TransAct (Miltenyi Biotec, #130-111-160) for 72h, then for exhausted cells they are re-stimulated every 48h. For chronic stimulation, cells were centrifuged, and the pellet was resuspended at 1-2×10^6^ cells/mL in fresh complete media containing TransAct (1:100) and IL-2 (100 U/mL). After the third stimulation, IL-15 (10 ng/mL, Miltenyi Biotec, #130-095-762) was added. Comparisons were made with CD8 T cells activated for 72h and then expanded for the duration of the experiment (expanded), and exhausted cells that were stimulated four times and then stained for TGlow at the same time as expanded cells.

### Automation

When possible, automation was performed with the Integra Bioscience AG systems: ViaFlow 96, ViaASSIST Plus with the D-One pipette and the 8-channel 5-125µL Voyager.

### TGlow

#### Cell seeding

Optical black 384-well plates (Perkin Elmer, #6057302) were coated with 0.01% poly-L-lysine (VWR International LTD, #P8920 or Merck, #A-00-5C) for two hours at room temperature or overnight at +4°C, washed twice with cell culture grade water (Corning, #15363651) and either used immediately or stored at +4°C for up to one week.

The samples were collected in a round-bottom 96-well plate and the cells were washed once in HBSS^+/+^ (Gibo^TM^, #14025050; 400g, 5 minutes), resuspended at 2-3.25×10^6^ cells/mL in HBSS^+/+^ and seeded onto the coated imaging plate at 30,000 – 45,000 cells per well (15µL). Each sample had 4-6 imaging replicates (one replicate= one well) randomly dispersed throughout the plate (excluding the edges), which also included isotype controls, single stained controls and no cell controls.

#### Cycle 1

Immediately after cell seeding, an equal volume of MitoTracker^TM^ Deep Red FM (Invitrogen^TM^, #M22426) was added to a final concentration of 0.16µM. The cells were incubated for 30 minutes at 37°C and fixed by adding paraformaldehyde (Thermo Scientific^TM^, #28906) to a final concentration of 3.7% for 20 minutes at room temperature. The fixative was washed away with three washes of 20µL HBSS^+/+^. The cells were permeabilized with 0.1% Triton X-100 (Sigma-Aldrich, #X100-100ML) for 10 minutes at room temperature and subsequently blocked with 1% bovine serum albumin (BSA, Sigma-Aldrich, #A5611) in HBSS^+/+^ for 30 minutes at room temperature.

After blocking, the cells were incubated on a rotator overnight at +4°C in a staining solution made in 1% BSA-HBSS^+/+^: CD25-FITC (1:100, clone M-A251, BioLegend^®^, #356116), Ki-67-FITC (1:100, clone Ki-67, BioLegend^®^, #350508), 120 nM Alexa Fluor™ 568 Phalloidin (Invitrogen^TM^, #A12380), and 2 µg/mL Hoechst 33342 (Invitrogen^TM^, #H3570). After overnight incubation and three washes of 20µL HBSS^+/+^, the cells were stored in 40µL HBSS^+/+^ to prevent evaporation. When possible, imaging of cycle 1 occurred within 48 hours of fixation.

#### Bleaching and Cycle 2

The fluorescent signals were removed by photobleaching (60 minutes of white light exposure followed by 30 minutes of UV light exposure) whilst the cells were in 40µL of a mild chemical bleaching buffer made of 24mM NaOH (stock: 1M, Sigma-Aldrich, #72068-100ML) and 4.5% H_2_O_2_ (stock: 30%, Sigma-Aldrich, #H1009-500ML) in PBS^+/+^. After bleaching, the cells were washed twice with HBSS^+/+^, blocked with 1% BSA in HBSS^+/+^, and stained overnight with the TGlow cycle 2 panel: 2.5µg/mL ConcanavalinA-Alexa Fluor™ 647 (Invitrogen™, #C21421), tubulin-PE (1:50, clone REA1136, Miltenyi Biotec, #130-119-540), 5µM BODIPY 505/15 (Invitrogen™ #D3921) and 2 µg/mL Hoechst 33342. The drug perturbation experiment had a different cycle 2 panel: 0.42µg/mL ConcanavalinA-Alexa Fluor™ 488 (Invitrogen™, #C11254), tubulin-PE (1:50, clone REA1136, Miltenyi Biotec, #130-119-540), and 2 µg/mL Hoechst 33342. After overnight incubation and three washes of 20µL HBSS^+/+^, the cells were stored in 40µL HBSS^+/+^ to prevent evaporation.

#### Microscopy

Cells were imaged using an Opera Phenix Plus High Content Screening System equipped with a PerkinElmer FLEX plate handling robot (PerkinElmer Inc.). Nine non-overlapping fields of view in a 3×3 grid were imaged per well. Images were acquired using a 40X water-immersion objective with 1 x 1 binning, yielding a pixel size of 0.149µm by 0.149µm. Sixteen focal planes were collected at 0.8µm spacing (total z-depth= 12.8µm), except for the drug dataset which used 0.5µm spacing, providing near-complete coverage of most activated cells. Having a complete 3D slice of the cells enables the use of deconvolution to account for artefacts introduced by the microscope’s point spread function (PSF) which greatly helps to improve contrast, segmentation and the ability to resolve fine details in the image, which is particularly important given the small size of lymphocytes (see ‘*Determination of the effective resolution’* below). The images were taken sequentially from green (excitation: 488 nm, emission filter: 522 nm bandwidth 25nm) to orange (excitation: 561 nm, emission filter: 599 nm bandwidth 30nm) to red (excitation: 640 nm, emission filter: 706 nm bandwidth 15nm), blue (excitation: 375 nm, emission filter: 456 nm bandwidth 23nm) and when appropriate, brightfield. Laser power and exposure times were calibrated on the controls in the first imaging plate, and then set constant for all subsequent plates. For cycle one the exposures in seconds and power (%) were as follows; 375: 0.06 (50%), 488: 0.3 (80%), 561: 0.1 (80%), 640: 0.12 (80%). For cycle two the exposures were as follows; 375: 0.06 (50%), 488: 0.04 (40%), 561: 0.1 (80%), 640: 0.1 (60%), and brightfield 0.2.

### Image processing

An in house image processing pipeline was developed to automate the process of data conversion, segmentation, registration, demultiplexing, illumination correction and feature extraction, with parallelization and efficient task execution using Nextflow *25.04.6* ^85^. First, raw image exports from PerkinElmer Harmony were converted to non-pyramidal CZYX OME-TIFF stacks ^86^, which resulted in one image stack per field. Each well was organized so it had a plate/row/col/field.ome.tiff folder structure. Channel names and pixel sizes were transferred from the Revity generated Index.xml to the appropriate metadata elements on the OME metadata (channel_names, physical_pixel_sizes). Each cycle of imaging is treated as a separate input plate in the raw data. This structure is suited for large datasets (tens of thousands of stacks) for use on filesystems that do not handle many small files, which was an issue with OME-ZARR and the raw exports from Harmony.

#### Flatfield correction

To correct for flatfields, A modified polynomial of a degree four in X and Y direction was fit to a randomly selected subset of images for a plate (Eq. 1).

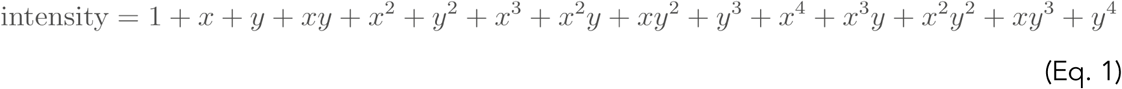

To address issues with sparse images we had in some datasets, 20 randomly selected images were first combined by max projection (compound image) prior to identifying the Otsu threshold, so a more constant threshold value was applied to each image. Images were thresholded using a two class Otsu threshold, which was then divided by a factor of four to have a lenient definition of the foreground signal. In total, 200 compound images were flattened into a single vector, removing background pixels, and combined with the other images. This was then used to fit the polynomial coordinates using ordinary least squares. To evaluate the fit, a second subset of random images was loaded and corrected for the learned flatfield, after which the image was binned into a 20×20 grid and an average foreground value calculated for each bin. This was used to evaluate the remaining relationship between the flatfield value and corrected image intensity. This is implemented in the tglow-pipeline script run_flatfield_estimation.py.

#### Deconvolution

When required, the images were deconvoluted using a GPU accelerated non-circulant version of the Richardson-Lucy algorithm (*CLIJ2-fft*, v0.27)^87^ with a point spread function (PSF) generated with the PS-Speck™ Microscope Point Source Kit (Invitrogen^TM^, #P7220) and imaged at 0.1nm intervals for 150 planes using the same plates, coating and objective as for the TGlow protocol. Nine fields were generated for each of the four channels.

PSFs were generated for each channel by first identifying spots in the PSF image (skimage blob_log), selecting spots with no close neighbours (20 pixels). The first spot was taken as the reference to which the other spots were aligned using phase cross correlation. Beads that registered poorly with the reference bead (>1.645 sd of the residual mean squared error) were removed. A common background value was calculated based on the edge of the stack, this was used to subtract from the average PSF in the next step. The bead images were averaged to derive the provisional PSF. The provisional PSF was then centred on its centre of mass in xyz, background value subtracted and divided by the maximum value to normalize it to 0-1, with any values falling below zero due to background subtraction being clipped to 0. The code is available in the tglow-pipeline repository in the script run_beads_to_psf.py.

To apply deconvolution for any given z-spacing the 0.1um PSF was matched to the dataset by sampling every n planes starting from the center depending on the z-spacing of the dataset. The non-circulant Richardson-Lucy algorithm was run with 100 iterations for each channel, keeping default values for the regularization parameter. Given we use 16bit-unit to store the data, but deconvolution converts the data into a 32bit float and can increase the intensity of pixels, we scale the deconvoluted images to a common max intensity of 5*65335 (value configurable in the pipeline) to ensure we do not clip the data and then save it to a 16 bit-unit OME-TIFF. The option is available to store the data 32 bit floats, but due to the number of images processed in this paper this was not feasible due to storage limitations. We note, once all deconvolutions are done, we scale each channel to a common optimal range based on the data (see *Batch effect scaling)*. This strategy ensures intensities remain comparable across images and are optimally used, as opposed to scaling to the max of the image, which would complicate interpretation of intensity information. The code to run the deconvolution is available in the tglow-pipeline script run_richardson_lucy.py.

#### Determination of the effective resolution

To determine the effective resolution pre and post deconvolution we calculated the full width half maximum (FWHM) of the bead images described above, before and after 100 interactions of the non-circulant Richardson-Lucy. For each channel, we selected all the bead crops, centered them on the center of mass. Then we filtered intensity outliers (MAD >3). The FWHM was then calculated by taking a center slice for each axis and fitting a gaussian distribution to the profile. The FWHM for each was taken as 2.35482 * sigma, where sigma is the standard deviation of the gaussian. We then filtered for poor fits of the gaussian by removing beads FWHM values >3 MADs. The mean over all remaining beads was taken as the final FWHM for that channel. This is implemented in the tglow-pipeline script run_measure_fwhm.py.

For the 375 channel; the mean FWHM were z=1.87μm y=0.43μm, and x=0.42μm before deconvolution, and z=0.66μm, y=0.24μm and x=0.24μm after. For the 488 channel; z=1.52μm, y=0.45μm, x=0.41μm before and z=0.49μm, y=0.24μm and x=0.22μm after. For the 561 channel; z=1.40μm, y=0.47μm and x=0.4μm before and z=0.49μm, y=0.23μm and x=0.22μm. Finally for the 640 channel; z=1.43μm, y=0.52μm and x=0.49μm before and z=0.51μm, y=0.24μm and x=0.23μm after deconvolution (**Suppl. Fig. 2**). These numbers represent the maximum achievable resolution with this microscope setup. We note that for these calculations we used the full stack of PSF beads spaced at 0.1μm to get the most accurate estimates of the theoretical maximum, but in the main acquisitions we use 0.8μm. However, as after deconvolution the z resolution is at most 0.66μm which falls below the plane spacing of 0.8μm we are limited by the plane spacing in the main dataset, so we put this as our maximum resolution post deconvolution.

#### Registration

To register image cycles, the DNA stain was used to calculate a translation matrix using the masked phase cross correlation (skimage^88^) on the foreground regions of the image (Otsu threshold). This gives a translation matrix for each of the images that was saved and used later on in the pipeline when combining the image stacks. During QC, the cell level Pearson correlation after registration between the two DNA channels was used to identify cells which registered well, as we observed that in some of our datasets, a portion of cells moved with respect to each other. We defined these badly registered cells by a Pearson R < 0.6 and they were discarded in downstream analysis. The code to run the registration is available in the tglow-pipeline script run_registration.py.

#### Segmentation

Segmentation is either run on the raw images when no deconvolution is chosen, or on the deconvoluted images when deconvolution is enabled. Images of nuclei and cells were segmented separately using the *Cellpose* V3 algorithm^15^ on the Hoechst and actin channels respectively, providing the nucleus channel to the cell segmentation for the purpose of declumping. The following non-default parameters were set: model=cyto2, cell size=75, nucleus size=60, cellprob_threshold –4 for the cell segmentation and 2 for the nucleus. In some cases the low cell density in some images meant the standard normalisation (normalised to 1-99th quantile) applied in *Cellpose* was not appropriate, as it clipped the signal values. Instead, images are pre-scaled to the 1-99.9th quantile and subsequently disabling scaling in *Cellpose*. To run *Cellpose*, GPU acceleration was enabled. The code to run cellpose is available in the tglow-pipeline script run_cellpose.py.

#### Batch effect scaling

To account for differences in average image intensities between batches of plates, we included a common control (Jurkat cells) that we expect to be relatively stable. We note that within an experiment, the microscope and staining settings are kept constant. To optimally use the dynamic range, we first perform a feature extraction on the unscaled data. This is used to determine the dynamic range of the data within actual cells, and the average differences between Jurkat intensities between plates. To ensure no artifacts are included in calculating these offsets, we filtered for cells which registered well (DNA-DNA R>0.6). These are used to determine the optimal scaling to fit the relevant data into a 16bit-uint and the offsets between average intensity in the Jurkats in each channel + plate. After deriving the scaling factors, the finalized images are produced, where each channel + plate is divided by its corresponding scaling factor. We note that scaling in this way can give rise to issues for background signals which remain constant between plates or batches, causing differences in minimal intensities observed in CellProfiler features. To alleviate this issue, we apply a ‘soft threshold’ or weight to the scaling factor based on intensity (Eq. 2). The weight for a pixel is determined by a sigmoid curve that ranges between 0 and 1 so it smoothly transitions the scale factor between background intensities and foreground intensities. Each plate gets its own sigmoid cure so the foreground and background regions are fairly determined if intensities vary between plates.

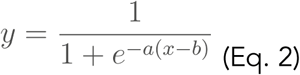

Where the slope a is defined as:

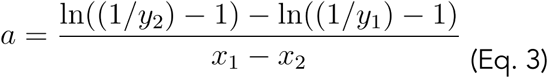

And the bias b is defined as:

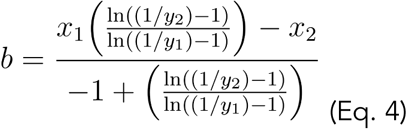

Where y1 and y2 are constants representing the desired output weights at intensity values x1 and x2 and are set to 1×10^−3^ and 1 – 1×10^−3^ respectively. For each plate and channel the value of x1 is determined by taking the median UpperQuartileIntensity of control images masked by the cell masks (e.g. the background intensity). The value for x2 is determined by taking the median Otsu threshold of control cells in the image. Finally, y represents the final output weight for the scaling factor and is bound between 0 and 1. In other words, the sigmoid ensures no scaling is done in background intensities, and then smoothly increases the scaling to regions of foreground intensity as defined by the Otsu threshold. This ensures that pixels with a value in the background region remain the same across batches, but the foreground signals that change due to staining differences or batch effects are scaled according to their appropriate scaling factor. This largely eliminated issues with min intensities associating to the scale factor, while correctly normalizing batch effect differences in the Jurkat controls.

In cases where only one plate is run in an experiment, the plate specific offset and sigmoid is omitted and just the dynamic range scaling is applied. We applied dynamic range scaling as we observed in some cases, low intensity channels would give rise to issues with texture measurements in CellProfiler, producing a lot of ties. The application of this dynamic range scaling to better fit the grey level co-occurance matrix in the 255 values CellProfiler uses, alleviated the problem. Scaling factors, biases and slopes were manually calculated in an R script on QCed cells, and supplied to subsequent runs of tglow-pipeline with the arguments rn_manualscale, rn_scale_bias, rn_scale_slope.

#### Finalizing images

The raw or deconvolved images, the flatfield corrections, scaling offsets, registration matrices and the segmentation masks were imported into a custom python script to stage them prior to running CellProfiler. Here the flatfield corrections are applied, any demultiplexing of nuclear stains is applied and files are saved in a CellProfiler compatible format for both 2D and 3D. For training DINO and visualizing the cells, the same images used as input for CellProfiler were taken and cell level crops were created based on the mask and saved in an HDF5 file where each group is a cell and each cell is a 4 dimensional matrix CZYX. The cell and nucleus masks are appended as the last 2 channels in the stack.

#### CellProfiler Feature extraction

Feature extraction was performed using CellProfiler *4.2.6* ^13^. ImageQuality metrics were calculated, and images with fewer than 5 cells were discarded. The primary objects were defined as cell (cell mask), nucleus (nucleus mask), cytoplasm (cell masks minus nucleus masks) and membrane (cell mask outline ± 10 pixels minus the nucleus region). The following features were extracted for each object and each channel: Radial Distribution, Texture, Intensity, Granularity, and AreaShape. Furthermore, a mitochondria mask was defined from the Mitotracker Deep-Red stain after removing the local background signal and applying a 2 class Otsu threshold. The binary mask was used to assign individual mito objects and a mitoNetwork mask (defined as all mito objects inside a single cell). This process results in 2,351 features which are trimmed to a standard number of 1,098 features by removing redundant and uninformative measurements (**Suppl. Tab. 1**). The nomenclature follows the following convention: *object_category_category-subtype_channel*, where object can be cell, cytoplasm (cyto), nucleus (nucl), membrane (memb) or mitochondria (mito, for the 18 features from mitochondria segmentation); category can be intensity, texture, granularity, radial distribution, correlation or area shape; and channel can be mito, er, dna, dna2, actin, cd25, ki67, lipids, tubulin). The texture features often include a series of numbers after the channel which describe their resolution, orientation and/or co-occurrence matrix scale.

#### DINO4Cells

As an alternative to the feature extraction from CellProfiler, DINO4Cells, a self-supervised vision transformer model, was employed to derive morphological embeddings of cell crops ^14^. The standard workflow was modified to enable extraction of single-channel embeddings, as in CellProfiler, reflecting the biological relevance of individual channels. Rather than learning a unified descriptor of cellular morphology across all stains simultaneously, the model was adapted to operate on a single stain at a time, enabling the extraction of feature representations for each channel independently. Since the modified model works on single-band images, the main difference from the original workflow is that individual channels are randomly sampled from each cell during training iterations. The model was trained on 239,805 cells derived from a CD4^+^ T cell time-course experiment (n= 193,756) and a CRISPR-perturbation experiment (n= 46,049), encompassing but not limited to the samples analyzed in this manuscript.

The model, a ViT-Base transformer with patch size 16, was therefore trained on a total of 1,918,440 images (239,805 cell crops × 8 channels) for 50 epochs, using a per-GPU batch size of 128. The learning rate was set to 5×10^−3^, and was linearly increased during the first 10 epochs as a warm-up strategy. All the default augmentations from the original DINO4Cells have been maintained, except for the addition of *RandomRotation* and the substitution of *RandomResizedCrop* with *RandomResizedCenterCrop*. At inference, features are extracted from each channel of each cell crop, resulting in a 8×768 matrix as a descriptor of single-cell morphology.

#### Image visualization

Images were visualized in *Napari* (v0.5.4, ^89^ or in R with *EBImage* (v4.48.0, ^90^. No contrast modifications were applied in any images shown in the manuscript. To ensure visibility, intensities for cells within a figure subpanel were normalized to the 99th quantile observed within that subset of cells for each channel separately.

### Feature analysis

The output of CellProfiler was imported into R and analysed with a set of functions that were packaged into the R package *tglow-r* which implements a Seurat like workflow for quality control and analysis of paired image level and single cell features.

#### Quality control

Quality control was performed sequentially at three levels: images, features and cells. Control groups were defined as sets of images, and their corresponding cells, that are expected to be biologically similar, such as knockout conditions, drug perturbations or cell-types.

First, images containing fewer than 5 or more than 1,000 cells were discarded. Next, outlier images were identified using the function *tglowr::find_outlier_pca*: for each control group, PCA was performed on CellProfiler-ImageQuality features. PCs explaining 75% of the variance within each control group were retained, and images were flagged as outliers based on their Euclidean distance in PCA space (FDR < 0.05).

Prior to feature quality control, poorly registered cells were removed by filtering out cells with a DNA-DNA Pearson correlation R < 0.6. Feature-level filtering was applied using a set of predefined criteria (**Suppl. Tab. 13**) to remove, for instance, blacklisted features^74^, redundant features, features with no unique values, features with many NA’s and features with low coefficient of variation. This is implemented in the functions *tglowr::calculate_feature_filters* and *tglowr::apply_feature_filters*.

Single cell quality control was first performed with hard thresholds to filter on cell size, number of nuclei, percentage of missing values, percentage of infinite values and number of neighbors (**Suppl. Tab. 14**). This is implemented in tglowr::calculate_object_filters and tglowr::apply_object_filters. Finally, cell outliers were identified either by applying *tglowr::apply find_outlier_pca* within each control group or during clustering of the full dataset in the PCA space. This QC enabled the removal of cells with fluorescent artifacts or poor segmentation, whilst keeping more rare populations like apoptotic cells in the dataset.

#### Feature transformation

To account for diverse distribution types, the features were transformed using a Box-Cox transformation^91^ which identifies the optimal exponent to approximate a normal distribution as closely as possible (*MASS::boxcox*)^92^. We note that this will not produce a normal distribution in all cases, just the best approximation that can be achieved by a power transform. As the BoxCox transformation accepts only positive values, we transform the data using equation 5 prior to running BoxCox transform.

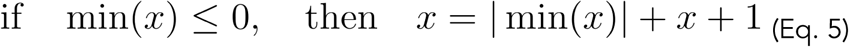

To find the optimal lambda, a range of values between –5 and 5 is tested in steps of 0.1. To improve speed, a random sample of 20,000 cells is taken to estimate the optimal lambda. The value which gives the maximum likelihood is chosen. A fudge factor is set to 0.2 which results in lambdas ranging between –0.2 and 0.2 having a log transform applied and values 0.8 and 1.2 having a lambda of 1 (no effect). This is referred to as the ‘transformed’ or ‘trans’ assay. This is implemented in the function *tglowr::apply_boxcox*. After transformation, all of the features were scaled to have a mean of 0 and standard deviation of 1, with the function *tglowr::scale_assay*.

#### Dimensionality reduction and clustering

Principal component analysis was performed using *irlba::prcomp_irlba*^93^, and clusters were identified using louvain clustering from *igraph*^94^ on an approximate kNN graph built using the implementation from *Seurat v5*^75^. The kNN was built using 30 PC’s and a k of 10. For each comparison done using this clustering strategy, multiple clustering resolutions were evaluated, after which one was chosen that best represented the major biological structures in the data. The results were subsequently embedded in a UMAP for visualisation using *uwot::umap*^95^. This workflow is implemented in *tglowr::calculate_pca*, *tglowr::calculate_umap* and *tglowr::calculate_clustering*.

#### Feature modelling

Single cell associations were performed using linear mixed models (lmerTest^96^ and lme4^97^. For CellProfiler features, these were performed on the scaled transformed assay. For DINO embeddings, associations were performed directly on the scaled principal components. The p-values for individual fixed effect coefficients were extracted using *lmerTest::lmer*. For other model comparisons with groups of coefficients (e.g. multiple doses or timepoints), a likelihood ratio test was performed using base R anova between a full and reduced model. The random effect structure included a nested random effect for ‘plate:well’ unless otherwise specified. This is implemented in *tglowr::calculate_lmm* and *tglowr::lmm_matrix*. In cases where donor or well means are modelled, no random effect structure is supplied and the ordinary least squares implementations *tglowr::calculate_lm* and *tglowr::lm_matrix* are used. Exact models used are described in the respective Supplementary Tables.

To identify marker features, the same base model (without an intercept) was used, however instead of testing if coefficients were zero, a contrast matrix was provided to lmerTest::contest to test if the class of interest differed significantly from the mean of the other classes, or compared to a reference class of interest. This is implemented in *tglowr::find_markers_lmm*. To use ordinary least squares instead of mixed models to find markers, set method=”lm”.

#### Identifying and reporting feature sets

Although individual features are modeled separately, the high redundancy among high-content imaging features makes it challenging to report associations, as highly correlated features often yield multiple redundant hits. To address this in reporting, we grouped similar features into “feature sets,” defined as clusters of highly correlated features. To do so, for each channel or channel category, we computed the feature–feature correlation matrix from the transformed assay, set negative correlations to 0, and derived a dissimilarity matrix as 1 – correlation. Complete linkage clustering was then applied, and the dendrogram was cut at a height of 0.3 (default, with cut-offs stable between 0 and 1, **Suppl. Fig. 4E**) to define feature sets. This is implemented in the function *tglowr::calculate_feature_clustering* and the results can be visualized with the function *tglowr::plot_markers*, where the percentage of significant feature sets instead of features are plotted. A feature set is considered significant if at least one feature within it shows an association; however, in practice, highly correlated features within a set typically move in the same direction and with the same magnitude and are summarized in the supplementary figures. Effect sizes are reported as the mean coefficient across features within each set. We note that CellProfiler features often have theoretical opposites, such as measures of pixel homogeneity versus heterogeneity, which are inherently anti-correlated. By only taking positive correlations forward we retain these features in separate sets, noting its impact and means and ranges, and refer to the supplementary tables for details.

Feature sets are named using the convention *channel_(category) cluster-number*, which is inherently non-descriptive. To aid interpretation, we typically report the results of the feature set, and in addition an illustrative feature from each set.

#### Assigning cell cycle phases

To assign cell-cycle phases, we plotted the raw integrated DNA intensity in the nucleus versus the log transformed integrated KI67 intensity in the nucleus. This gives a similar plot to common assignments of cell cycle in flow cytometry^98^. Gates were set manually in each dataset to identify the major populations of G0, G1, S, G2M. Differences in cell-cycle composition across conditions were assessed using linear models applied to the mean percentage of cells in each phase per donor.

#### Differential abundance testing

To test for enrichment of variables in neighbourhoods we applied Milo^99^. Milo was applied using default settings, on PCA’s derived from either DINO embeddings or the transformed assay. We input 30 PCs for each of these tests. To link neighbourhood enrichments to discrete clusters we assigned neighborhoods to their most common discrete cluster, and the average effect size of neighborhoods and their significance was evaluated. As an alternative strategy, we calculated the cell proportions for each donor in the dataset and evaluated the differences in proportion using linear regression. Alternatively, differences in cluster composition across conditions were assessed using linear models applied to the mean percentage of cells in each cluster per donor.

#### Variance Decomposition

To obtain estimates of overall explained variance per variable we applied the variance partitioning analysis which uses mixed linear models to estimate the contribution of variables to the overall variance in the feature space^100^. We used either the scaled transformed CellProfiler features or the raw DINO embeddings. In this framework, all categorical variables are modelled as random effects. We applied this to both single cell level data and well level means.

#### Selection of cells to show as representative images

Representative cells were identified in one of two ways depending if the feature of interest is categorical (e.g. a cluster label) or continuous. For categorical features, we calculate 30 principal components on the scaled assay data (either DINO or CellProfiler based). For each group in the feature we subset the PCA space to include only cells in that group, after which we calculate the squared euclidean distance to the centroid of the group. This is done by centering each PC in the subset to its mean, and subsequently calculating the sum of squares over the 30 PCs for each cell. We then pick the cell with the lowest distance as the “index cell” for that group. To find its neighbours we find its k nearest neighbours using a k nearest neighbours graph in PCA space using RANN:nn2^101^. This is implemented in tglowr::fetch_representative_object_nn. For continuous features we select the feature of interest, and calculate the 5th, 50th and 95th quantiles (unless otherwise specified) and take these cells that have these values as “index cells”. For each index cell we then take the k cells upstream and downstream of the index cell to extend the set. This is implemented in tglowr::fetch_representative_object_quantiles.

## Code & data availability

The code for the tglow-pipline, tglow-r, adaptation of DINO4cells and analysis scripts are available on the TrynkaLab github. The pipeline and R package have additional documentation on their wiki pages. The tglow-core package supporting the pipeline is available on PyPi as well.

– TGlow R package: https://github.com/TrynkaLab/tglow-r
– TGlow pipeline: https://github.com/TrynkaLab/tglow-pipeline
– TGlow core: https://github.com/TrynkaLab/tglow-core
– DINO-TGlow: https://github.com/TrynkaLab/tglow-dino4cells

Raw and processed images as well as segmentation masks & metadata have been deposited in the EBI bioimage archive^102,103^ and will be made public upon publication. Imaging features and model weights have been deposited in EBI BioStudies^102^. The data used in this study has been deposited under the following accessions:

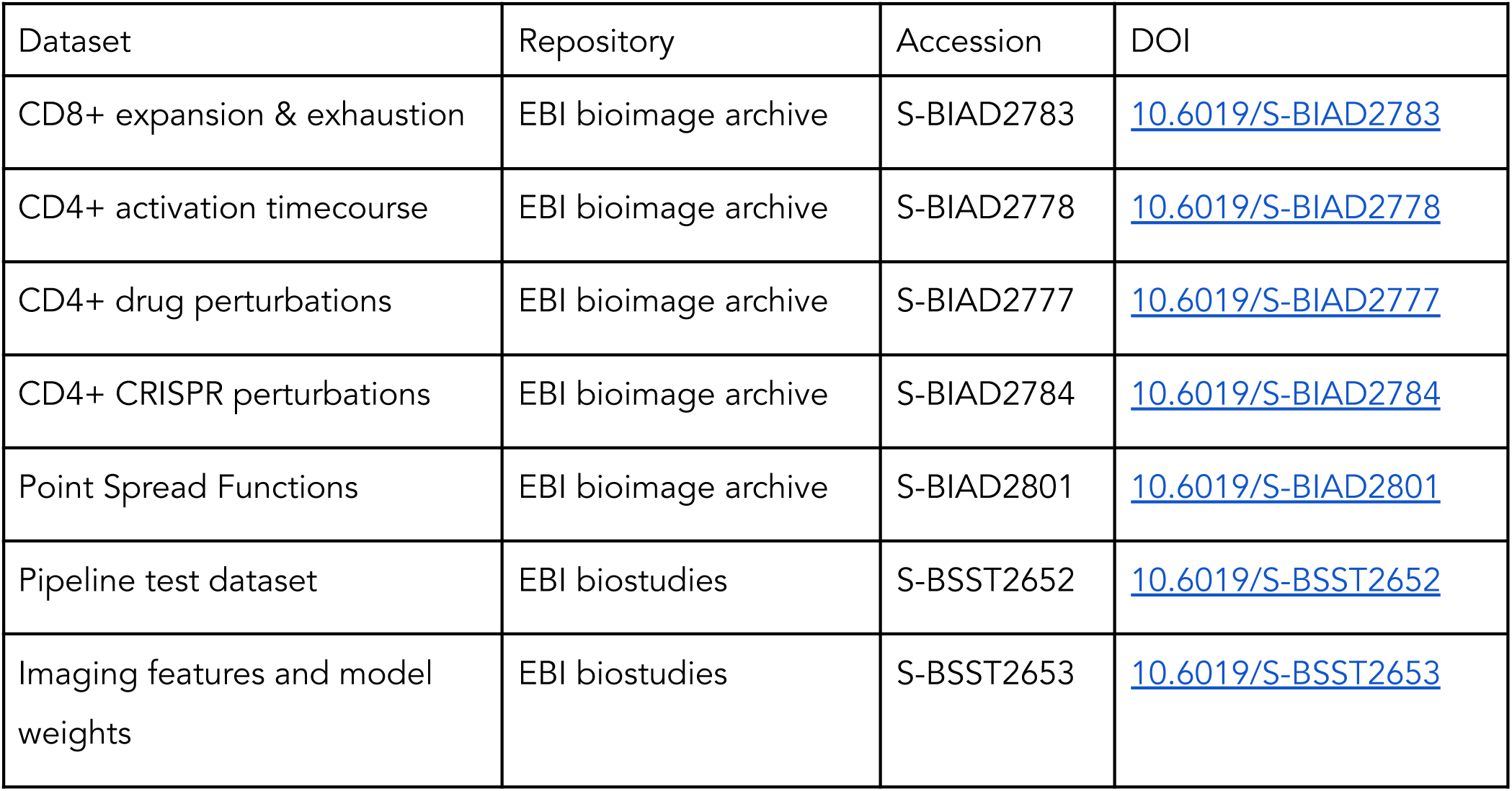

## Acknowledgements

We thank the Wellcome Sanger Institute Human Genetics Informatics department for their assistance in setting up the backup infrastructure used for imaging data. We thank the Cellular Genetics Informatics department (Tong Li and Martin Prete) for their assistance in streamlining the workflow from the Phenix microscope to the high performance compute infrastructure as well as providing input into the best practices for the pipeline. We thank Kwasi KawaKwa for his continued assistance in running and maintaining the Phenix microscope. We thank Brian Northan for developing the deconvolution package CLIJ2-fft and helping to debug several technical issues, without which the pipeline would not have been possible. We thank Filip Konopacki for his generosity in sharing his timeslots on the Phenix microscope. We thank Cristina Cotobal Martin, Laura Richardson and May Hu for their help with sample processing. We thank Leopold Parts for his insightful comments and input throughout the project. We thank Gareth Griffiths for his help in validation of the editing efficiencies. The authors wish to acknowledge the support of the Cytometry Core Facility at Wellcome Sanger Institute.

## Author contributions

J.C.M. was involved in: Formal analysis, Conceptualization, Funding acquisition, Methodology, Investigation, Visualization, Writing – original draft; O.B.B. was involved in: Formal analysis, Conceptualization, Funding acquisition, Methodology, Visualization, Writing – original draft, Software; M.A.O. was involved in: Formal analysis, Investigation, Methodology; F.C. was involved in: Formal analysis, Software; A.J.M.R. was involved in: Investigation; A.H. was involved in: Methodology; F.L. was involved in: Investigation; T.L. was involved in: Software, Resources; K.K was involved in: Resources; A.O.S was involved in: Methodology; C.A.G was involved in: Supervision; O.B. was involved in: Resources; C.P.J. was involved in: Investigation, Writing – review & editing; M.C. was involved in: Conceptualization, Writing – review & editing; G.T. was involved in: Conceptualization, Funding acquisition, Supervision, Writing – original draft.

All roles defined according to the Credit taxonomy: https://credit.niso.org/

## Supplementary Materials

### List of Supplementary Figures

● Suppl. Fig. 1 – Evaluation of impact of scaling on Jurkat cells in the drug dataset.
● Suppl. Fig. 2 – Effective resolution estimates based on bead images.
● Suppl. Fig. 3 – Schematic outlining the TGlow image processing workflow
● Suppl. Fig. 4 – Supplementary data for Cell Profiler features.
● Suppl. Fig. 5 – Supplementary data for the drug perturbation experiment.
● Suppl. Fig. 6 – Supplementary profiles for the drug perturbation experiment.
● Suppl. Fig. 7 – Supplementary data for the CD4^+^ T cell activation time course.
● Suppl. Fig. 8 – Supplementary data for the CRISPR perturbation dataset.
● Suppl. Fig. 9 – Supplementary data for the CD8^+^ T cell exhaustion dataset.

### List of Supplementary Tables

● Suppl. Tab. 1 – List of CellProfiler features
● Suppl. Tab. 2 – Feature set membership for each experiment
● Suppl. Tab. 3 – Association statistics from linear mixed-effects models fitted to CellProfiler features in the drug dataset
● Suppl. Tab. 4 – Cluster marker CellProfiler features for mitochondria clustering in the drug dataset
● Suppl. Tab. 5 – Association statistics from linear mixed-effects models fitted to DINO-derived principal components in the time course dataset
● Suppl. Tab. 6 – Cluster marker CellProfiler features for merged-DINO clustering in the timecourse dataset
● Suppl. Tab. 7 – Association statistics from linear mixed-effects models fitted to Cell Profiler features in the time course dataset
● Suppl. Tab. 8 – Association statistics from linear mixed-effects models fitted to DINO derived principal components (PCs) in the CRISPR dataset
● Suppl. Tab. 9 – Association statistics from linear mixed-effects models fitted to Cell Profiler features in the CRISPR dataset
● Suppl. Tab. 10 – Association statistics from linear mixed-effects models fitted to Cell Profiler features in the exhaustion dataset
● Suppl. Tab. 11 – Cluster marker CellProfiler features for CellProfiler clustering in the exhaustion dataset
● Suppl. Tab. 12 – Guide sequences and editing efficiencies for the knockouts used in the crispr dataset.
● Suppl. Tab. 13 – Quality control filter table applied to features
● Suppl. Tab. 14 – Quality control filter table applied to cells.

