## Supplementary Figures for "Multiplexed high-content imaging uncovers morphological diversity of lymphocyte activation and dysfunction"

### Supplementary Materials

**List of Supplementary Figures**


- Suppl. Fig. 1 – Evaluation of impact of scaling on Jurkat cells in the drug dataset.
- Suppl. Fig. 2 – Effective resolution estimates based on bead images.
- Suppl. Fig. 3 – Schematic outlining the TGlow image processing workflow
- Suppl. Fig. 4 – Supplementary data for Cell Profiler features.
- Suppl. Fig. 5 – Supplementary data for the drug perturbation experiment.
- Suppl. Fig. 6 – Supplementary profiles for the drug perturbation experiment.
- Suppl. Fig. 7 – Supplementary data for the $CD4^+$ T cell activation time course.
- Suppl. Fig. 8 – Supplementary data for the CRISPR perturbation dataset.
- Suppl. Fig. 9 - Supplementary data for the $CD8^+$ T cell exhaustion dataset.


**List of Supplementary Tables**


- Suppl. Tab. 1 – List of CellProfiler features
- Suppl. Tab. 2 – Feature set membership for each experiment
- Suppl. Tab. 3 – Association statistics from linear mixed-effects models fitted to CellProfiler features in the drug dataset
- Suppl. Tab. 4 – Cluster marker CellProfiler features for mitochondria clustering in the drug dataset
- Suppl. Tab. 5 – Association statistics from linear mixed-effects models fitted to DINO-derived principal components in the time course dataset
- Suppl. Tab. 6 – Cluster marker CellProfiler features for merged-DINO clustering in the timecourse dataset
- Suppl. Tab. 7 – Association statistics from linear mixed-effects models fitted to Cell Profiler features in the time course dataset
- Suppl. Tab. 8 – Association statistics from linear mixed-effects models fitted to DINO derived principal components (PCs) in the CRISPR dataset
- Suppl. Tab. 9 – Association statistics from linear mixed-effects models fitted to Cell Profiler features in the CRISPR dataset
- Suppl. Tab. 10 – Association statistics from linear mixed-effects models fitted to Cell Profiler features in the exhaustion dataset
- Suppl. Tab. 11 – Cluster marker CellProfiler features for CellProfiler clustering in the exhaustion dataset
- Suppl. Tab. 12 – Guide sequences and editing efficiencies for the knockouts used in the crispr dataset.
- Suppl. Tab. 13 – Quality control filter table applied to features
- Suppl. Tab. 14 – Quality control filter table applied to cells.

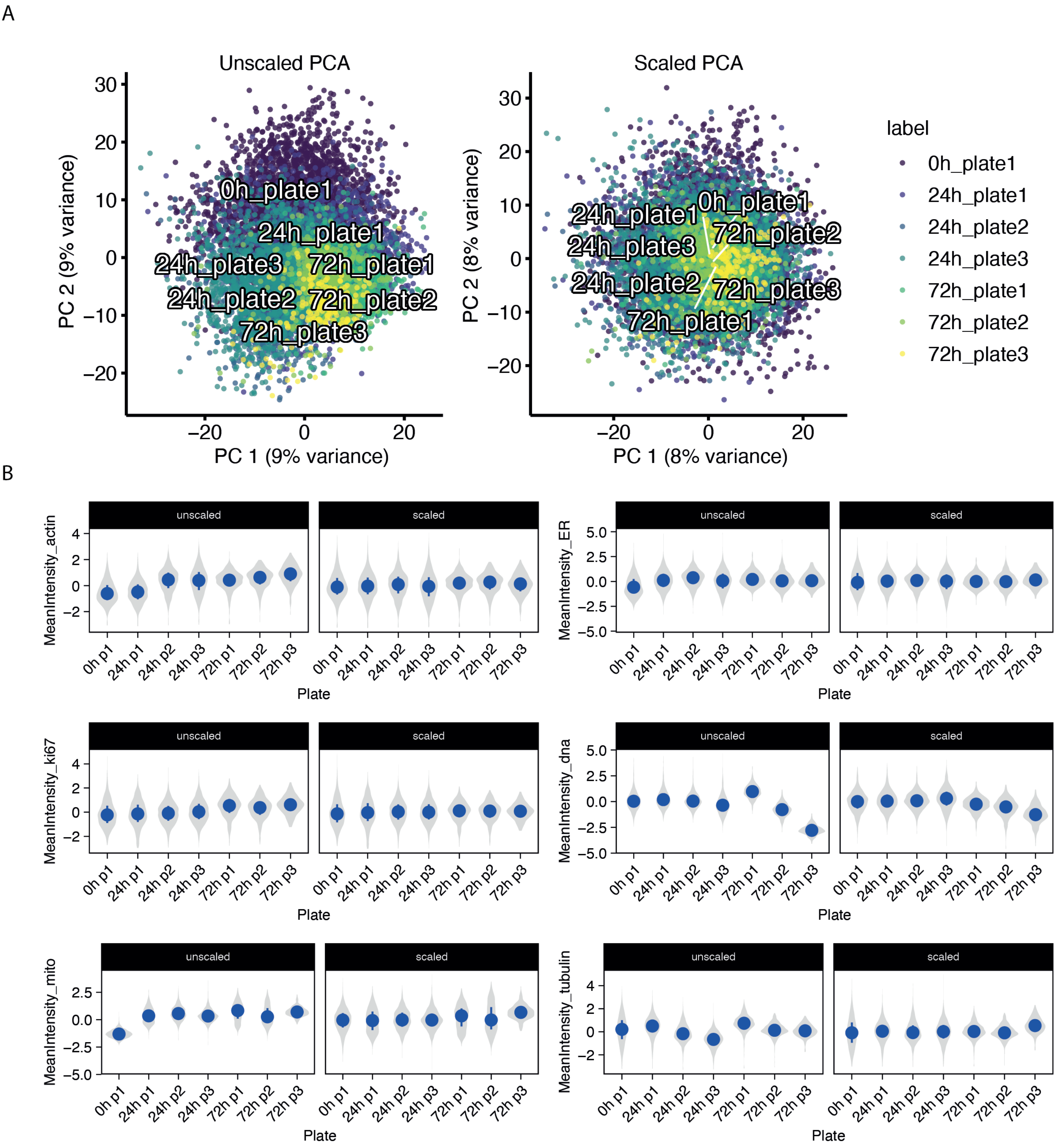
A
Unscaled PCA
Scaled PCA
PC 1 (9% variance)
PC 2 (9% variance)
PC 1 (8% variance)
PC 2 (8% variance)
label
0h_plate1
24h_plate1
24h_plate2
24h_plate3
72h_plate1
72h_plate2
72h_plate3
B
unscaled
scaled
MeanIntensity_actin
MeanIntensity_ER
MeanIntensity_ki67
MeanIntensity_dna
MeanIntensity_mito
MeanIntensity_tubulin
0h p1
24h p1
24h p2
24h p3
72h p1
72h p2
72h p3
Plate

**Suppl.Fig. 1 - Evaluation of impact of scaling on Jurkat cells in the drug dataset. A)** Dot plot of principal components one and two of the full feature space in the Jurkat cells before (left) and after (right) scaling each plate using the strategy described in the **Methods**. Each dot is colored by its corresponding combination of timepoint (0h, 24h, 72h) and plate (p1, p2, p3). The centroid of each plate is labelled in white. **B)** Cell-level transformed mean intensity per scaled channel (facet) in the Jurkat cells before (unscaled, left) and after (scaled, right) scaling. The y-axis shows scaled feature values, x-axis shows the plate.

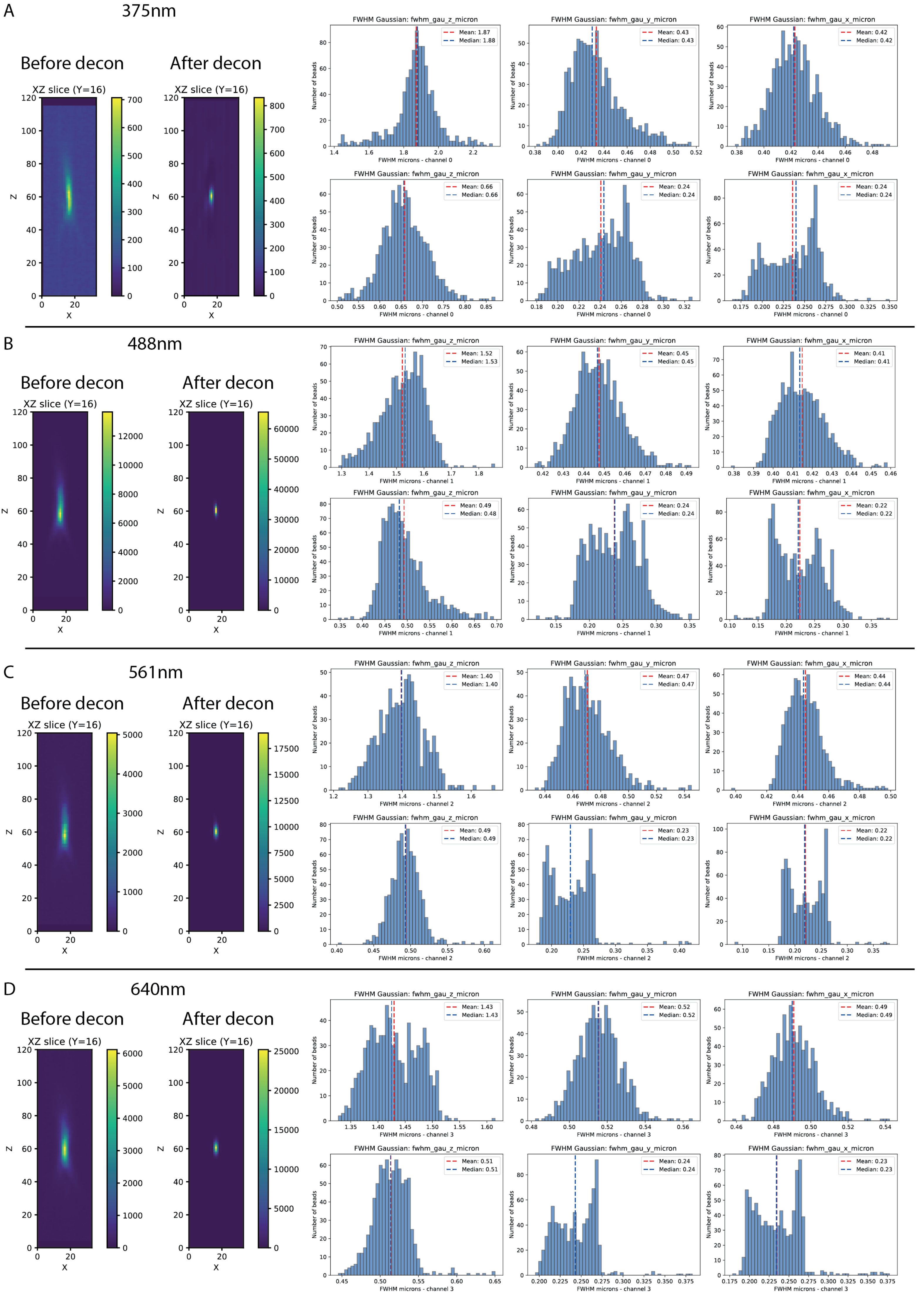

A
375nm
Before decon
After decon
XZ slice (Y=16)
XZ slice (Y=16)
FWHM Gaussian: fwhm_gau_z_micron
Mean: 1.87
Median: 1.88
FWHM microns - channel 0
FWHM Gaussian: fwhm_gau_y_micron
Mean: 0.43
Median: 0.43
FWHM microns - channel 0
FWHM Gaussian: fwhm_gau_x_micron
Mean: 0.42
Median: 0.42
FWHM microns - channel 0
FWHM Gaussian: fwhm_gau_z_micron
Mean: 0.66
Median: 0.66
FWHM microns - channel 0
FWHM Gaussian: fwhm_gau_y_micron
Mean: 0.24
Median: 0.24
FWHM microns - channel 0
FWHM Gaussian: fwhm_gau_x_micron
Mean: 0.24
Median: 0.24
FWHM microns - channel 0
Number of beads
B
488nm
Before decon
After decon
XZ slice (Y=16)
XZ slice (Y=16)
FWHM Gaussian: fwhm_gau_z_micron
Mean: 1.52
Median: 1.53
FWHM microns - channel 1
FWHM Gaussian: fwhm_gau_y_micron
Mean: 0.45
Median: 0.45
FWHM microns - channel 1
FWHM Gaussian: fwhm_gau_x_micron
Mean: 0.41
Median: 0.41
FWHM microns - channel 1
FWHM Gaussian: fwhm_gau_z_micron
Mean: 0.49
Median: 0.48
FWHM microns - channel 1
FWHM Gaussian: fwhm_gau_y_micron
Mean: 0.24
Median: 0.24
FWHM microns - channel 1
FWHM Gaussian: fwhm_gau_x_micron
Mean: 0.22
Median: 0.22
FWHM microns - channel 1
C
561nm
Before decon
After decon
XZ slice (Y=16)
XZ slice (Y=16)
FWHM Gaussian: fwhm_gau_z_micron
Mean: 1.40
Median: 1.40
FWHM microns - channel 2
FWHM Gaussian: fwhm_gau_y_micron
Mean: 0.47
Median: 0.47
FWHM microns - channel 2
FWHM Gaussian: fwhm_gau_x_micron
Mean: 0.44
Median: 0.44
FWHM microns - channel 2
FWHM Gaussian: fwhm_gau_z_micron
Mean: 0.49
Median: 0.49
FWHM microns - channel 2
FWHM Gaussian: fwhm_gau_y_micron
Mean: 0.23
Median: 0.23
FWHM microns - channel 2
FWHM Gaussian: fwhm_gau_x_micron
Mean: 0.22
Median: 0.22
FWHM microns - channel 2
D
640nm
Before decon
After decon
XZ slice (Y=16)
XZ slice (Y=16)
FWHM Gaussian: fwhm_gau_z_micron
Mean: 1.43
Median: 1.43
FWHM microns - channel 3
FWHM Gaussian: fwhm_gau_y_micron
Mean: 0.52
Median: 0.52
FWHM microns - channel 3
FWHM Gaussian: fwhm_gau_x_micron
Mean: 0.49
Median: 0.49
FWHM microns - channel 3
FWHM Gaussian: fwhm_gau_z_micron
Mean: 0.51
Median: 0.51
FWHM microns - channel 3
FWHM Gaussian: fwhm_gau_y_micron
Mean: 0.24
Median: 0.24
FWHM microns - channel 3
FWHM Gaussian: fwhm_gau_x_micron
Mean: 0.23
Median: 0.23
FWHM microns - channel 3

**Suppl.Fig. 2 - Effective resolution estimates based on bead images.** For each channel we calculated the full width at half maximum (**Methods**) for each bead in two fields of view before and after deconvoluting the images with the derived PSF. **A)** Left to right: Center slice of a randomly selected bead before deconvolution in Z and X direction, pixel sizes are 0.1µm in Z and 0.149µm in X. Next to it a center slice in Z and X of separate randomly selected bead after deconvolution. The histograms show the full width at half maximum estimates in Z,Y,X (columns) over all beads, with the top row of 3 prior to deconvolution, and the bottom row post-deconvolution. **B)** As A but for the 488nm channel. **C)** As A but for the 561m channel. **D)** As A but for the 640nm channel (edited)

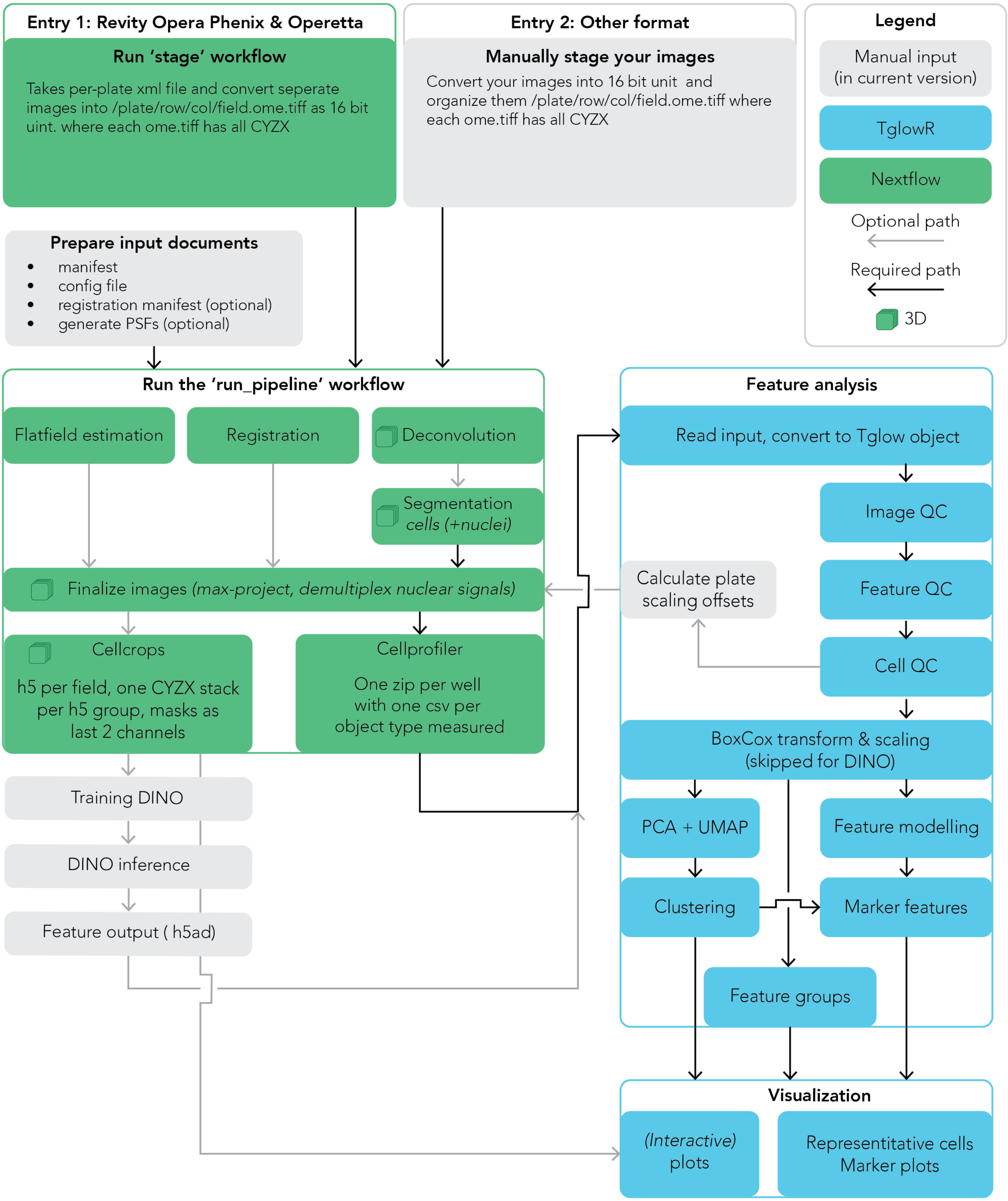

Entry 1: Revity Opera Phenix & Operetta
Run 'stage' workflow
Takes per-plate xml file and convert seperate images into /plate/row/col/field.ome.tiff as 16 bit uint. where each ome.tiff has all CYZX
Entry 2: Other format
Manually stage your images
Convert your images into 16 bit unit and organize them /plate/row/col/field.ome.tiff where each ome.tiff has all CYZX
Legend
Manual input (in current version)
TglowR
Nextflow
Optional path
Required path
3D
Prepare input documents
manifest
config file
registration manifest (optional)
generate PSFs (optional)
Run the 'run_pipeline' workflow
Flatfield estimation
Registration
Deconvolution
Segmentation cells (+nuclei)
Finalize images (max-project, demultiplex nuclear signals)
Cellcrops
h5 per field, one CYZX stack per h5 group, masks as last 2 channels
Cellprofiler
One zip per well with one csv per object type measured
Training DINO
DINO inference
Feature output ( h5ad)
Feature analysis
Read input, convert to Tglow object
Image QC
Calculate plate scaling offsets
Feature QC
Cell QC
BoxCox transform & scaling (skipped for DINO)
PCA + UMAP
Feature modelling
Clustering
Marker features
Feature groups
Visualization
(Interactive) plots
Representitative cells Marker plots

**Suppl.Fig. 3 - Schematic outlining the TGlow image processing workflow.** Grey boxes indicate a step in the process that requires manual input. Green boxes are implemented in Nextflow through the tglow-pipeline. Blue boxes are steps implemented in the tglow-r package. Grey arrows indicate steps that are optional to get results. Black arrows show steps that are required.

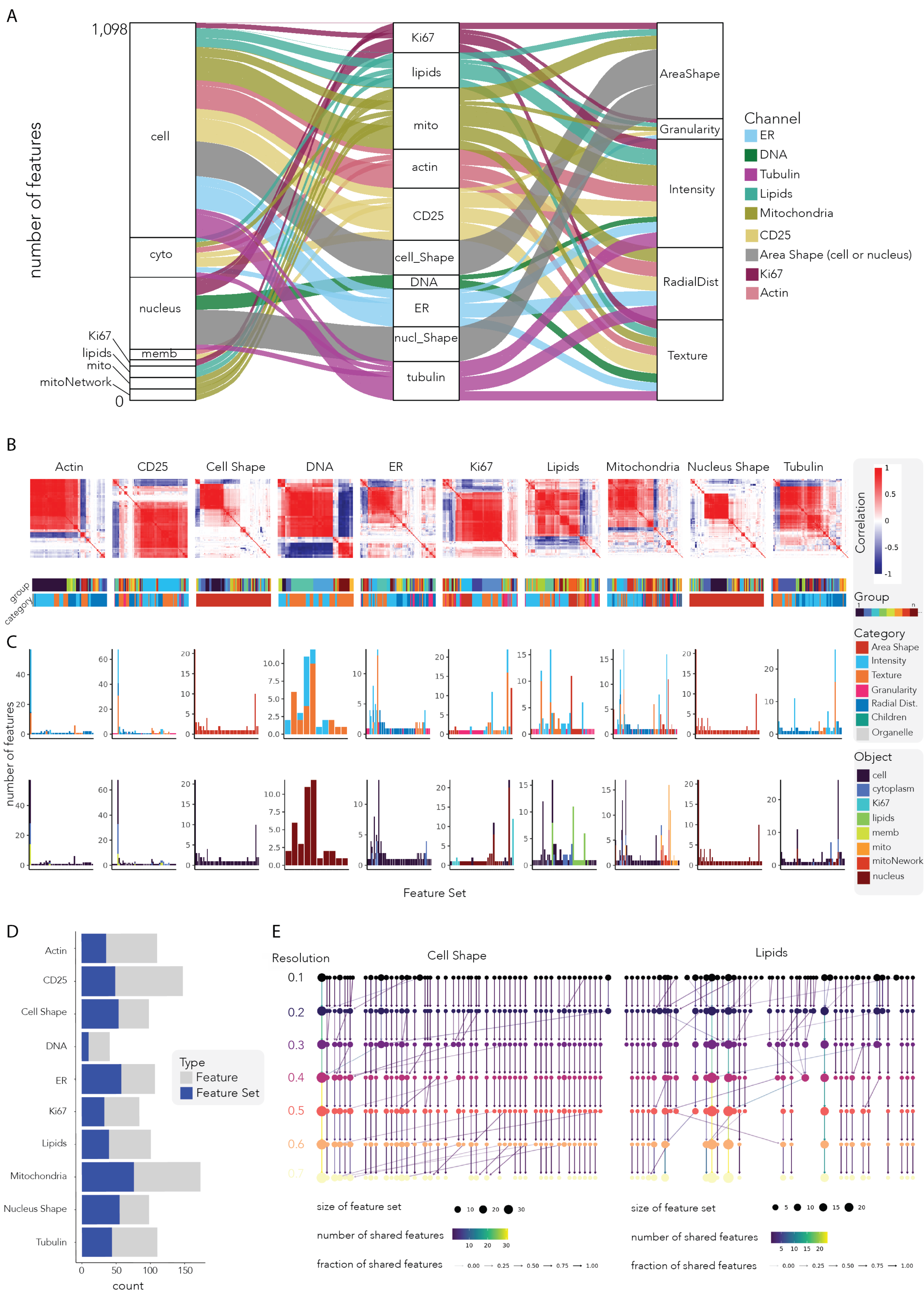

A
B
C
D
E
number of features
cell
cyto
nucleus
memb
Ki67
lipids
mito
mitoNetwork
Ki67
lipids
mito
actin
CD25
cell_Shape
DNA
ER
nucl_Shape
tubulin
AreaShape
Granularity
Intensity
RadialDist
Texture
Channel
ER
DNA
Tubulin
Lipids
Mitochondria
CD25
Area Shape (cell or nucleus)
Ki67
Actin
Actin
CD25
Cell Shape
DNA
ER
Ki67
Lipids
Mitochondria
Nucleus Shape
Tubulin
Correlation
group
category
Group
Category
Area Shape
Intensity
Texture
Granularity
Radial Dist.
Children
Organelle
Object
cell
cytoplasm
Ki67
lipids
memb
mito
mitoNework
nucleus
number of features
Feature Set
Type
Feature
Feature Set
count
Resolution
Cell Shape
Lipids
size of feature set
number of shared features
fraction of shared features

**Suppl.Fig. 4 - Supplementary data for Cell Profiler features. A)** Alluvial plot displaying the standard number of features after feature quality control (n = 1,098) and the relationship between the compartments (cell, cytoplasm (cyto), membrane (memb), nucleus, Ki67, lipids, mitochondria (mito) and mitoNetwork), channels (Ki67, lipids, mito, actin, CD25, cell_Shape, DNA, ER, nucleus shape (nucl_Shape) and tubulin) and measurements (AreaShape, Granularity, Intensity, RadialDistribution (RadialDist) and Texture). Colors map to the channels: ER: light blue, DNA: green, tubulin: purple, lipids: teal, mitochondria: green, CD25: yellow, AreaShape: dark blue, Ki67: burgundy. **B-E)** Data displayed is of the $CD4^+$ T cell activation dataset. **B)** Representative heatmap of the feature-feature correlation per channel (-1: blue, 0: white, +1: red), Above are displayed the feature category: Children (teal), AreaShape (red), Granularity (pink), Intensity (light blue), RadialDistribution (dark blue; RadialDist), Texture (orange) and Organelle (grey; amalgamation of mito, mitoNetwork, Ki67 and lipid objects) and designated feature set (arbitrary multi-colored) at a treeheight of 0.3. **C)** Barplot showing the number of features (y-axis) per feature set (x-axis), faceted by channel and coloured by category (above; Children (teal), AreaShape (red), Granularity (pink), Intensity (light blue), RadialDistribution (dark blue; RadialDist), Texture (orange) and Organelle (grey; amalgamation of mito, mitoNetwork, Ki67 and lipid objects) or object (below; Cell (dark blue), cytoplasm (medium blue), Ki67 (light blue), lipids (green), membrane (light green), mito (orange; mitochondria), mitoNetwork (red) and nucleus (brown). **D)** Barplot showing the number of features (grey) and number of feature sets (blue) per channel at a dissimilarity correlation of 0.3. **E)** Clustree visualization of two representative stains (Cell Shape, left and Lipids, right) showing the relationship between feature sets across increasing resolution parameters. Each node corresponds to a feature set, with node size proportional to the number of features in that set and node color corresponding to resolution. Edges indicate feature transitions between clusters at adjacent resolutions, with edge color and with proportional to the number and fraction of shared features, respectively.

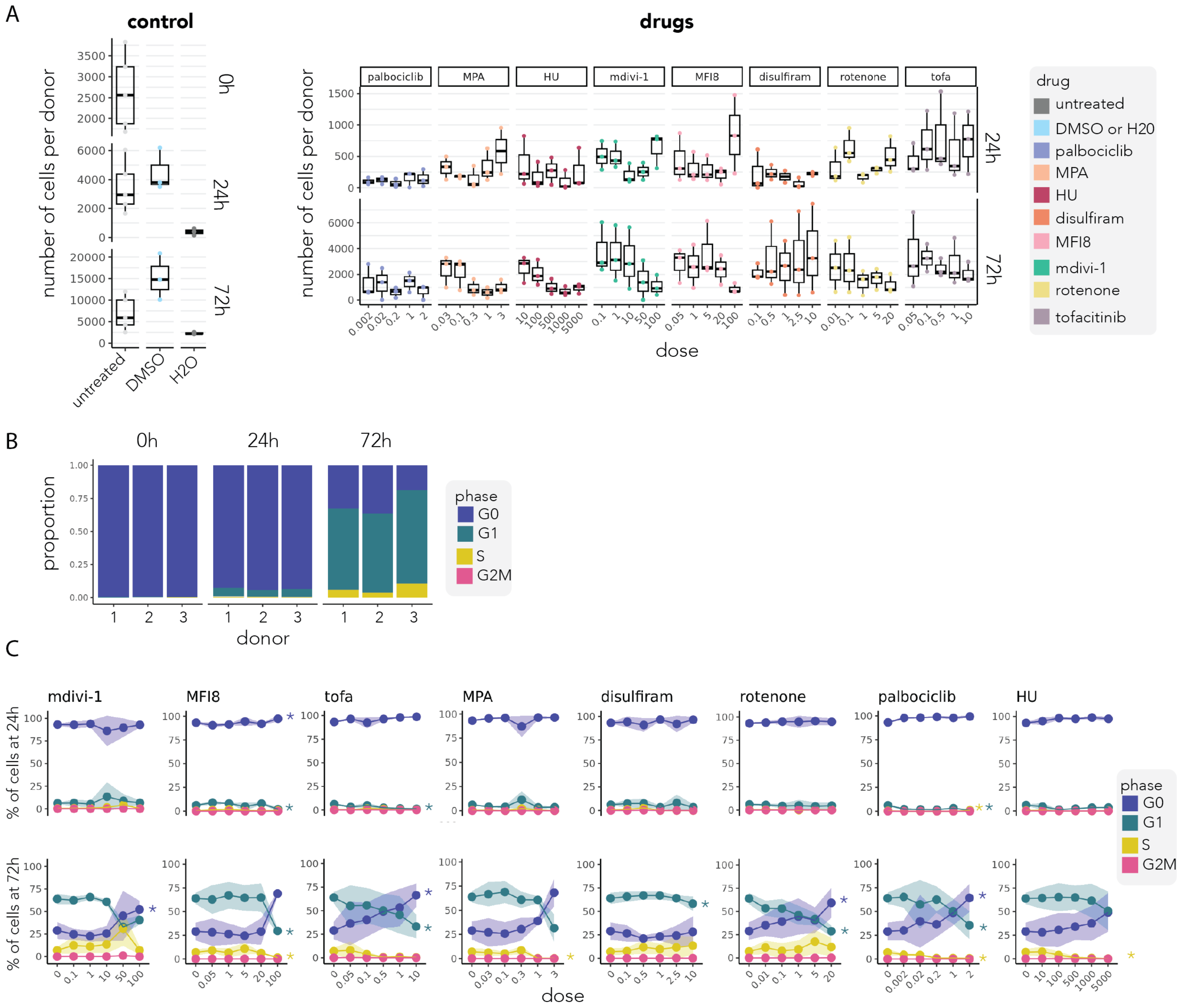
A
control
drugs
number of cells per donor
0h
24h
72h
untreated
DMSO
H2O
palbociclib
MPA
HU
mdivi-1
MFI8
disulfiram
rotenone
tofa
dose
drug
untreated
DMSO or H20
palbociclib
MPA
HU
disulfiram
MFI8
mdivi-1
rotenone
tofacitinib
B
proportion
donor
phase
G0
G1
S
G2M
C
mdivi-1
MFI8
tofa
MPA
disulfiram
rotenone
palbociclib
HU
% of cells at 24h
% of cells at 72h
dose

**Suppl.Fig. 5 - Supplementary data for the drug perturbation experiment. A)** Barplot, overlaid with a dotplot, showing the number of cells per donor for the controls (untreated, DMSO and H2O; left) at 0h, 24h and 72h and for the compounds at 24h and 72h (right). The dots are colored by treatment: untreated (grey), DMSO or H2O (blue), palbociclib (purple), MPA (peach), HU (red), disulfiram (orange), MFI8 (pink), mdivi-1 (green), rotenone (yellow) and tofacitinib (grey-purple). **B)** Proportion plot showing the fraction of cells of the DMSO control in each cell cycle phase per donor and per timepoint (0h, 24h and 72h): G0 (blue), G1 (green), S (yellow) and G2M (pink). **C)** For each drug (column) and both timepoints (24h and 72h, rows), the fraction of cells in each cell cycle phase per donor, shown as mean (dot) and SD (ribbon); G0 (blue), G1 (green), S (yellow), G2M (pink). Asterisk (*) indicate significant dose effects (model $p<= 0.05$).

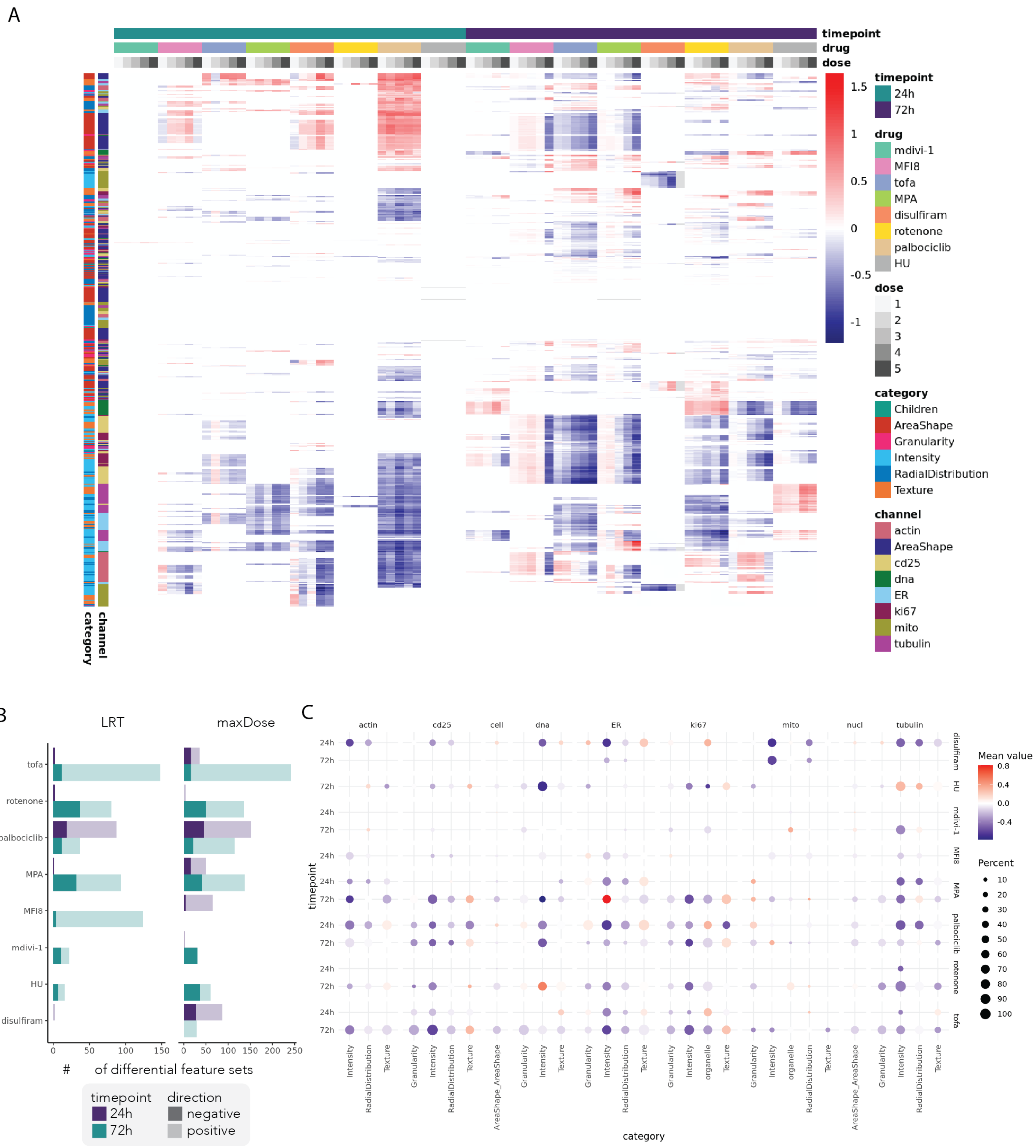

A
timepoint
drug
dose
timepoint
24h
72h
drug
mdivi-1
MFI8
tofa
MPA
disulfiram
rotenone
palbociclib
HU
dose
1
2
3
4
5
category
Children
AreaShape
Granularity
Intensity
RadialDistribution
Texture
channel
actin
AreaShape
cd25
dna
ER
ki67
mito
tubulin
1.5
1
0.5
0
-0.5
-1
category
channel
B
LRT
maxDose
tofa
rotenone
palbociclib
MPA
MFI8
mdivi-1
HU
disulfiram
drug
0 50 100 150
0 50 100 150 200 250
### of differential feature sets
timepoint
24h
72h
direction
negative
positive
C
actin
cd25
cell
dna
ER
ki67
mito
nucl
tubulin
timepoint
24h
72h
disulfiram
HU
mdivi-1
MFI8
MPA
palbociclib
rotenone
tofa
Intensity
RadialDistribution
Texture
Granularity
AreaShape_AreaShape
organelle
category
Mean value
0.8
0.4
0.0
-0.4
Percent
10
20
30
40
50
60
70
80
90
100

**Suppl.Fig. 6 - Supplementary profiles for the drug perturbation experiment. A)** Heatmap of the coefficients of all significant features (n= 972) for each timepoint_drug_dose from the model *feature ~ dose_factor + donor + (1 | plate:well)*. Values of insignificant feature sets (by LRT and maxDose) are set to 0. Coefficients range from -1.5 (blue) to 0 (white) to 1.5 (red). Samples are annotated by timepoint (24h: teal, 72h: purple), drug (palbociclib: purple, MPA:peach, HU: red, disulfiram: orange, MFI8: pink, mdivi-1: green, rotenone: yellow and tofacitinib: grey-purple) and dose (1 to 5, in increasing shades of grey). Features are annotated by channel (ER: light blue, DNA: green, tubulin: purple, lipids: teal, mitochondria: green, CD25: yellow, AreaShape: dark blue, Ki67: burgundy) and category (AreaShape: red, Granularity: pink, Intensity: light blue, RadialDistribution: dark blue, Texture: orange). **B)** Barplot depicting the number of differential feature sets per drug and timepoint (24h in purple, 72h in blue) modelled as *feature ~ dose_factor + donor + (1 | plate:well)* and averaged per feature set, with significance determined by LRT (left) or maximum dose (maxDose, right. The shading indicates the number of feature sets that have a positive (light) or negative (dark) effect size. **C)** Dot plot summarizing significant (maxDose) feature sets by drug/timepoint (rows) and channel (columns, n = 9). Dot size denotes the proportion of significant features in that channel; color encodes mean effect size: -0.8 (blue) to 0 (white) and +0.8 (red).

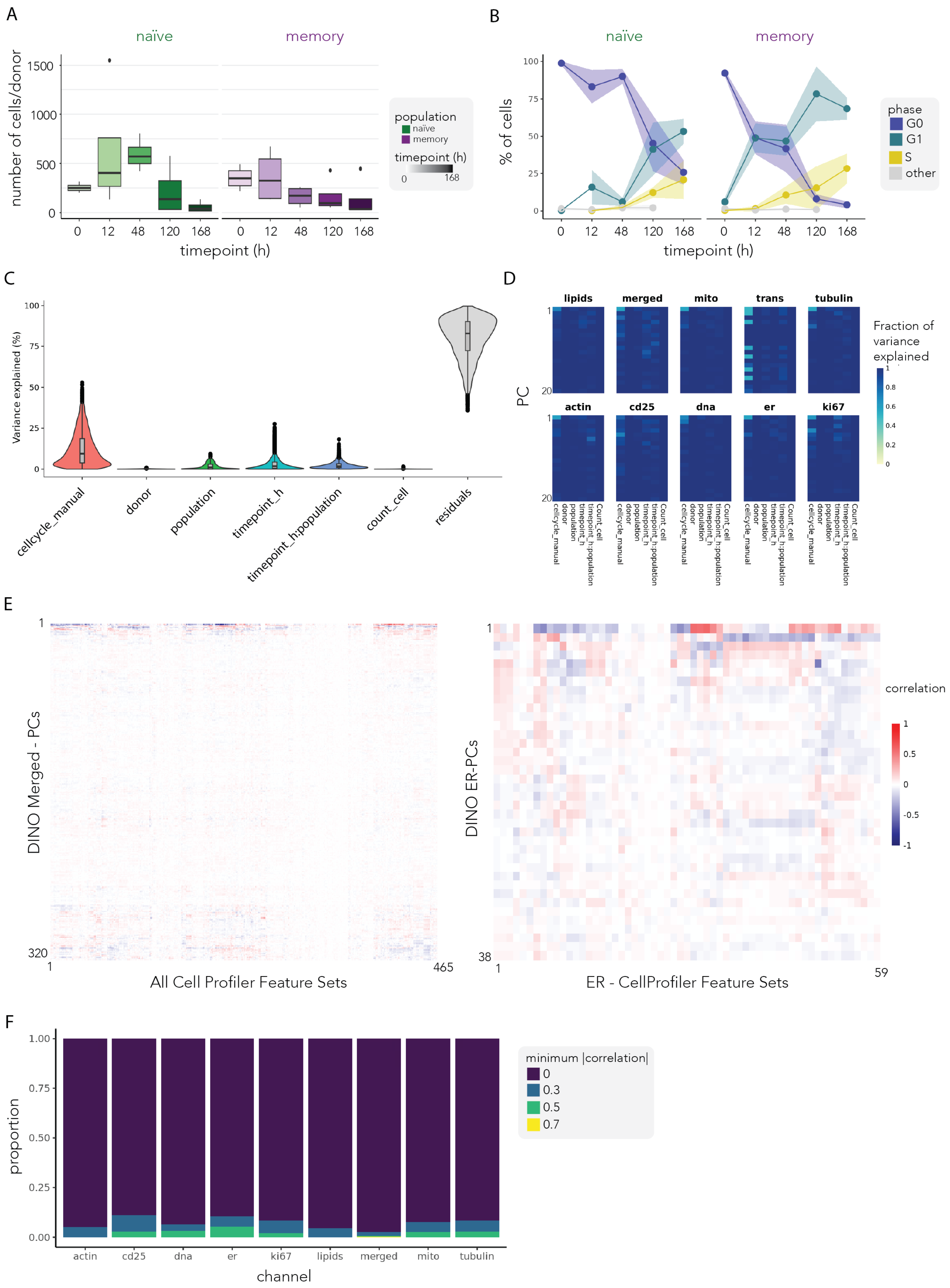
A
naïve
memory
number of cells/donor
1500
1000
500
0
0
12
48
120
168
timepoint (h)
population
naïve
memory
timepoint (h)
0
168
B
naïve
memory
% of cells
100
75
50
25
0
timepoint (h)
phase
G0
G1
S
other
C
Variance explained (%)
100
75
50
25
0
cellcycle_manual
donor
population
timepoint_h
timepoint_h:population
count_cell
residuals
D
lipids
merged
mito
trans
tubulin
actin
cd25
dna
er
ki67
PC
1
20
Fraction of variance explained
1
0.8
0.6
0.4
0.2
0
cellcycle_manual
donor
population
timepoint_h
timepoint_h:population
Count_cell
E
DINO Merged - PCs
1
320
465
All Cell Profiler Feature Sets
DINO ER-PCs
38
59
ER - CellProfiler Feature Sets
correlation
1
0.5
0
-0.5
-1
F
proportion
1.00
0.75
0.50
0.25
0.00
actin
cd25
dna
er
ki67
lipids
merged
mito
tubulin
channel
minimum |correlation|
0
0.3
0.5
0.7

**Suppl.Fig. 7 - Supplementary data for the CD4$^{+}$ T cell activation timecourse. A)** Barplot showing the number of cells per donor for the naïve (green; left) and memory (purple; right) populations and timepoints (x-axis; increasing darkness for increasing time). **B)** The fraction of cells in each cell cycle phase per donor, shown as mean (dot) and SD (ribbon); G0 (blue), G1 (green), S (yellow) and other (grey). **C)** Violin plot overlaid by a boxplot summarising, for each embedding of the DINO-merged representation (n = 6,144), the proportion of variance of each, explained by each variable and the residuals after having accounted for all other variables. The variancePartition R package was used to model *embedding ~ population + timepoint_h + population:timepoint_h + cellcycle_manual + donor + Count_cell* using (Methods). **D)** Heatmap showing the fraction of variance explained (0-1; yellow-dark blue) by the factors (population, timepoint_h, population:timepoint_h, cellcycle_manual, donor, Count_cell; column) in the model PC ~ factor. The PCs (rows) were calculated per channel (column facet) and only the first 20 PCs were used. **E)** Heatmap showing the correlation (-1:blue, 0: white, +1: red) between all CellProfiler features sets (x-axis, n = 465) and all DINO merged-PCs (y-axis, n = 320) and all ER CellProfiler feature sets (x-axis, n = 59) and all DINO ER-PCs (y-axis, n = 38). **F)** Proportion plot showing the fraction of embeddings per channel which have a minimum absolute correlation to a CellProfiler feature of 0.7 (yellow, 1 embeddings), 0.5 (green, 8 embeddings), 0.3 (blue, 23 embeddings) or 0 (purple).

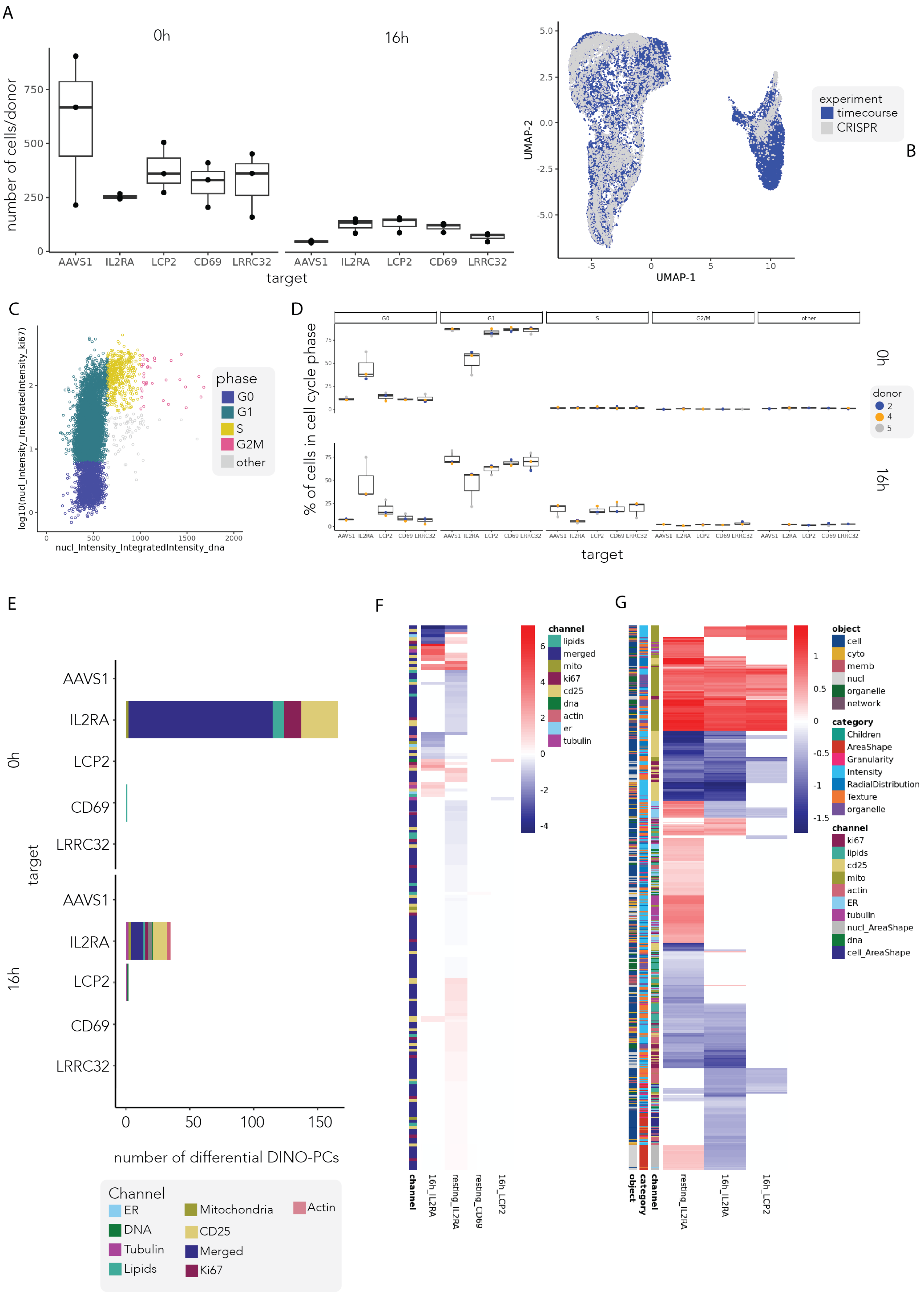
A
0h
16h
number of cells/donor
target
AAVS1
IL2RA
LCP2
CD69
LRRC32
B
UMAP-1
UMAP-2
experiment
timecourse
CRISPR
C
phase
G0
G1
S
G2M
other
log10(nucl_Intensity_IntegratedIntensity_ki67)
nucl_Intensity_IntegratedIntensity_dna
D
G0
G1
S
G2/M
other
% of cells in cell cycle phase
donor
2
4
5
E
number of differential DINO-PCs
Channel
ER
DNA
Tubulin
Lipids
Mitochondria
CD25
Merged
Ki67
Actin
F
channel
lipids
merged
mito
ki67
cd25
dna
actin
er
tubulin
16h_IL2RA
resting_IL2RA
resting_CD69
16h_LCP2
G
object
cell
cyto
memb
nucl
organelle
network
category
Children
AreaShape
Granularity
Intensity
RadialDistribution
Texture
organelle
channel
ki67
lipids
cd25
mito
actin
ER
tubulin
nucl_AreaShape
dna
cell_AreaShape
resting_IL2RA
16h_IL2RA
16h_LCP2

**Suppl.Fig. 8 - Supplementary data for the CRISPR perturbation dataset. A)** Barplot overlaid by a dotplot showing the number of cells per donor each target (x-axis) per timepoint (0h:left and 16h: right). **B)** UMAP embedding from the DINO-merged assay where each cell is colored by its experiment origin: $CD4^{+}$ T cell activation time course (blue) or CRISPR perturbation screen (grey). **C)** Cell cycle dot plot showing log10 (nuclear Ki67 integrated intensity) versus raw nuclear DNA integrated intensity and coded by cell cycle phase: G0 (blue), G1 (green), S (yellow), G2/M (pink) and other (grey). **D)** For each timepoint (0h, 16h) and perturbation, barplot overlaid by a dotplot showing the percentage of cells in each cell cycle phase per donor, with dots representing donors (grey, orange, blue). **E)** barplot showing the number of significant differential DINO-PCs per perturbation and timepoint, colored by channel: actin (pink), CD25 (yellow), DNA (green), Ki67 (burgundy), lipids (teal), merged (dark blue), mito (olive green) and tubulin (purple). **F)** Heatmap of the coefficients of all significant DINO-PCs (n= 123) for each timepoint_perturbation from the model *PC ~ target + timepoint + target:timepoint + donor + (1 | plate:well)*. Coefficients range from -4 (blue) to 0 (white) to 6 (red). DINO-PCs are annotated by channel (ER: light blue, DNA: green, tubulin: purple, lipids: teal, mitochondria: green, CD25: yellow, merged: dark blue, Ki67: burgundy). **G)** Heatmap of the coefficients of all significant features (n= 721) for each timepoint_perturbation from the model *PC ~ target + timepoint + target:timepoint + donor + (1 | plate:well)*. Coefficients range from -1.5 (blue) to 0 (white) to 1.5 (red). Features are annotated by channel (ER: light blue, DNA: green, tubulin: purple, lipids: teal, mitochondria: green, CD25: yellow, AreaShape: dark blue, Ki67: burgundy), category (AreaShape: red, Granularity: pink, Intensity: light blue, RadialDistribution: dark blue, Texture: orange) and object (cell: darkblue, cytoplasm: yellow, membrane: pink, nucleus (nucl): grey, organelle: green and network: dark grey).

A

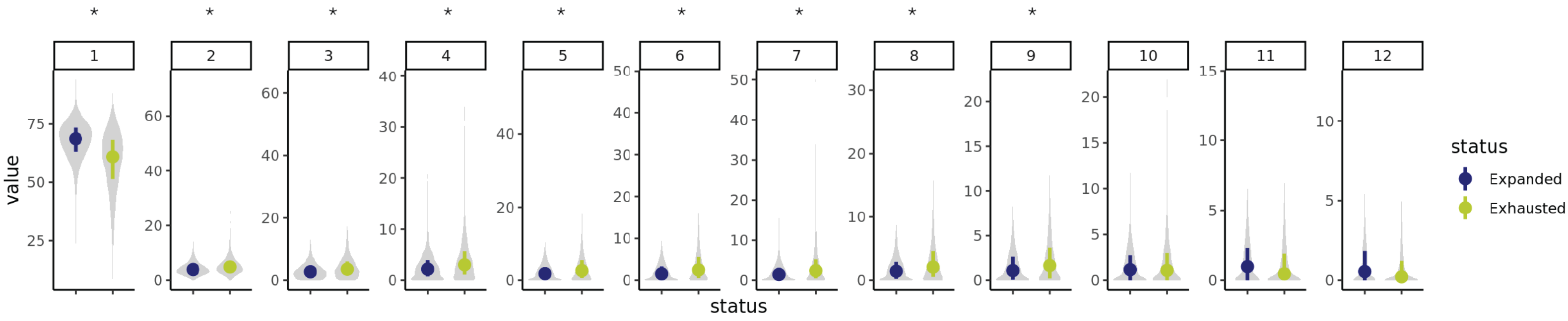
*
1
2
3
4
5
6
7
8
9
10
11
12
value
status
status
Expanded
Exhausted

B

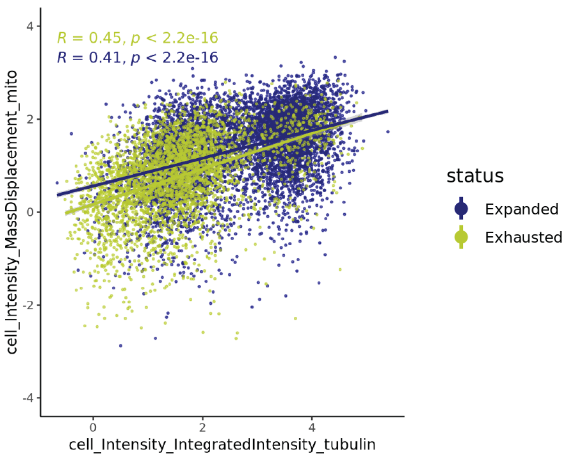
R = 0.45, p < 2.2e-16
R = 0.41, p < 2.2e-16
cell_Intensity_MassDisplacement_mito
cell_Intensity_IntegratedIntensity_tubulin
status
Expanded
Exhausted

**Suppl.Fig. 9 - Supplementary data for the CD8$^+$ T cell exhaustion dataset. A)** Single-cell violin plots of the raw lipid granularity spectra (1 to 12, faceted), with the median and interquantile range for each status indicated in a coloured point and line: expanded (blue) and exhausted (green). Significant differences ($p_{adj} \leq 0.05$ of the LRT) are shown with an asterisk (*). **B)** Dotplot of the transformed assay measurements of tubulin_Intensity_IntegratedIntensity (x-axis) and cell_Intensity_MassDisplacement_mito (y-axis). Each cell is colored by status: expanded (blue) and exhausted (green).
